# A morphology and secretome map of pyroptosis

**DOI:** 10.1101/2024.04.26.591386

**Authors:** Michael J. Lippincott, Jenna Tomkinson, Dave Bunten, Milad Mohammadi, Johanna Kastl, Johannes Knop, Ralf Schwandner, Jiamin Huang, Grant Ongo, Nathaniel Robichaud, Milad Dagher, Andrés Mansilla-Soto, Cynthia Saravia-Estrada, Masafumi Tsuboi, Carla Basualto-Alarcón, Gregory P. Way

**Affiliations:** Department of Biomedical Informatics, University of Colorado School of Medicine, USA; Assay.Works GmbH, Regensburg, Germany; Nomic Bio, Montreal, Québec, Canada; Department of Chemistry and Biotechnology, University of Tokyo, Tokyo, Japan; Health Sciences Department, University of Aysén, Coyhaique, Chile; Anatomy and Legal Medicine Department, University of Chile, Santiago, Chile

## Abstract

Pyroptosis represents one type of Programmed Cell Death (PCD). It is a form of inflammatory cell death that is canonically defined by caspase-1 cleavage and Gasdermin-mediated membrane pore formation. Caspase-1 initiates the inflammatory response (through IL-1β processing), and the N-terminal cleaved fragment of Gasdermin D polymerizes at the cell periphery forming pores to secrete pro-inflammatory markers. Cell morphology also changes in pyroptosis, with nuclear condensation and membrane rupture. However, recent research challenges canon, revealing a more complex secretome and morphological response in pyroptosis, including overlapping molecular characterization with other forms of cell death, such as apoptosis. Here, we take a multimodal, systems biology approach to characterize pyroptosis. We treated human Peripheral Blood Mononuclear Cells (PBMCs) with 36 different combinations of stimuli to induce pyroptosis or apoptosis. We applied both secretome profiling (nELISA) and high-content fluorescence microscopy (Cell Painting). To differentiate apoptotic, pyroptotic and control cells, we used canonical secretome markers and modified our Cell Painting assay to mark the N-terminus of Gasdermin-D. We trained hundreds of machine learning (ML) models to reveal intricate morphology signatures of pyroptosis that implicate changes across many different organelles and predict levels of many pro-inflammatory markers. Overall, our analysis provides a detailed map of pyroptosis which includes overlapping and distinct connections with apoptosis revealed through a mechanistic link between cell morphology and cell secretome.

## Introduction

Cell death pathways such as apoptosis and pyroptosis are defined canonically via cleaved caspases and specific morphologies.^1–4^ Pyroptosis (originally discovered in macrophages) is caspase-1 dependent and forms membrane pores, resulting in nuclear compaction and DNA fragmentation.^5^ These membrane pores form through Gasdermin D cleavage and translocation of its N-terminal domain to the cell and mitochondrial plasma membranes, where subsequent pores enable the secretion of pro-inflammatory markers.^6^ Conversely, apoptosis, one form of canonically non-inflammatory cell death, is caspase-3 dependent and causes membrane blebbing, nuclear compaction, and DNA fragmentation.^7^ Though defined separately from pyroptosis, recent research suggests they may have important overlaps. For example, both caspase-1 and caspase-3, in certain cases, are involved in both apoptosis and pyroptosis.^8,9^

With a growing appreciation for the complex intersection of cell death processes^10,11^, we sought to acquire and analyze comprehensive systems representations of the secretome and cell morphology to disambiguate apoptosis from pyroptosis. Furthermore, we focused on pyroptosis because several links between pyroptosis and disease have emerged in recent years. Aberrant regulation of pyroptosis has been implicated in inflammatory disease, cancer, chemotherapy resistance, and autoimmunity.^12,13^ Therefore, manipulating pyroptosis with therapeutic agents represents a new, underexplored frontier for many diseases. Unfortunately, incomplete knowledge of pyroptosis, and cell death single-molecule pleiotropy poses decreases the pace of therapeutic development. To develop new therapeutic agents targetting pyroptosis, the full landscape of the cellular pyroptotic states need to be elucidated and differentiated from other forms of cell death. We begin charting a map of pyroptosis using unbiased systems biology readouts of cell morphology and cell secretome. This will map pyroptosis as a process rather than as an individual caspase or other molecule in isolation.

Systems biology provides tools to measure comprehensive representations of biological processes by measuring holistic features in the same cell.^14,15^ There are many different ways to measure systems biology, each providing complementary views into cell function. Scientists have used systems biology to measure cell death complexities. For example, Sato et al. measured gene expression of cells undergoing necrosis (another form of cell death), and apoptosis to understand how genes differentially regulate these pathways.^16^ Further, Yu et al. discovered a gene expression signature of pyroptosis in low-grade glioma patients.^17^ In general, emerging assays and new approaches for studying complex diseases have enabled more extensive systems biology characterization of cell death.^18–21^ While recent work has characterized disease-specific transcriptome differences^22–24^, pyroptosis has been sparsely characterized via other systems biology views including high-content imaging and secretome^25^.

High-content microscopy is an emerging systems biology measurement. Microscopy yields robust information that is not captured by single molecule approaches such as sequencing, or the human eye.^26,27^ One of the most common high-content microscopy assays is Cell Painting. Cell Painting fluorescently stains the cell’s nucleus, mitochondria, endoplasmic reticulum (ER), cytoplasmic RNA, nucleoli, actin, Golgi apparatus, and plasma membrane.^28^ From these images, scientists extract high-content representations using a bioinformatics approach known as image-based profiling.^29^ These high-content representations have mostly been used for drug discovery^30,31^, but recent research has applied this approach for basic research studies, such as predicting cell health, toxicity, and cancer cell resistance.^31–35^ While the most common image-based profiling approach is to aggregate single-cell measurements per well, single-cell approaches enable more systematic analyses and are becoming increasingly common.^36^ For example, Schorpp et al. developed a machine-learning approach to predict apoptosis and necrosis from Cell Painting data at single-cell resolution.^37^ Similar advances in ELISA technologies have enabled the simultaneous acquisition of hundreds of secreted markers to form systems biology representations of the secretome, and recent technology allows the simultaneous measurement of secretome and cell morphology from the same wells.^38,39^

In this study, we applied Cell Painting and nELISA (Nomic Bio, 187 secretome markers) to profile cell morphology and secretome representations of pyroptosis and apoptosis. We modified the Cell Painting assay, swapping the cytoplasmic RNA stain with a marker for cleaved N-terminal Gasdermin D protein, which is the canonical marker of pyroptosis pore formation.^6,40^ We applied this assay to human Peripheral Blood Mononuclear Cells (PBMCs) perturbed with 36 chemical perturbations at different doses that induced or inhibited either apoptosis or pyroptosis (e.g., lipopolysaccharide [LPS], nigericin, flagellin, hydrogen peroxide [H202], etc.). We confirmed pyroptosis and apoptosis activity through the presence of canonical secreted markers, but individual pyroptosis treatments induced a complex secretome landscape. We also conducted extensive comparative analysis through training hundreds of machine-learning models to uncover distinctive cell morphology patterns associated with pyroptosis. Using cell morphology data, we successfully predicted levels of biologically significant pro-inflammatory cytokines such as IL-1β and TNF-α. We discovered key differences in nuclei, actin, Golgi apparatus, and plasma membrane morphology and Gasdermin D distribution. Pyroptotic cells showed different morphologies than apoptotic cells across all organelles we marked. Deep learning networks of single-cell morphology profiles accurately distinguished pyroptotic and apoptotic cells. In summary, our analysis revealed both shared and unique responses in pyroptosis and apoptosis and established a mechanistic link between cell morphology and the cell secretome.

## Results

### Experimental design and cell death process validation

We induced either apoptosis or pyroptosis by treating human PBMCs with 36 different combinations of pyroptosis and apoptosis-inducing and inhibiting agents (**Supplemental Table 1**). To isolate pyroptosis, we treated cells with LPS, flagellin, or a mixture of LPS and nigericin treated simultaneously.^41–47^ To isolate apoptosis, we treated cells with thapsigargin, which is an ER stress-mediated apoptotic inducer.^45,48–51^ To induce generic cell death we treated cells with H2O2. In addition, we also incubated some wells with apoptotic and pyroptotic inhibitors prior to treatment. We used disulfiram to inhibit Gasdermin D-mediated pore formation^52–55^ and Z-VAD-FMK to inhibit the proteolytic activity of caspases.^43,46,49,52^ Combining inhibitors with inducers maximizes our ability to isolate specific cell death pathways.

We incubated cells for one hour with an inhibitor or control, followed by six hours of cell death induction stimuli at varying dosages. We chose six hours for our assay time point to profile cells undergoing cell death rather than profiling cells that were already dead (**Supplemental Figure 1**). We included six to eight replicate wells of each inhibition/induction combination (36 combinations) within one 384-well plate. For a complete plate map, see **Supplemental Figure 2**. We applied a modified Cell Painting assay paired with secretome profiling to measure two systems biology representations of human PBMCs treated with this specific combination of apoptotic and pyroptotic agents (**Figure 1**).

**Figure 1.**
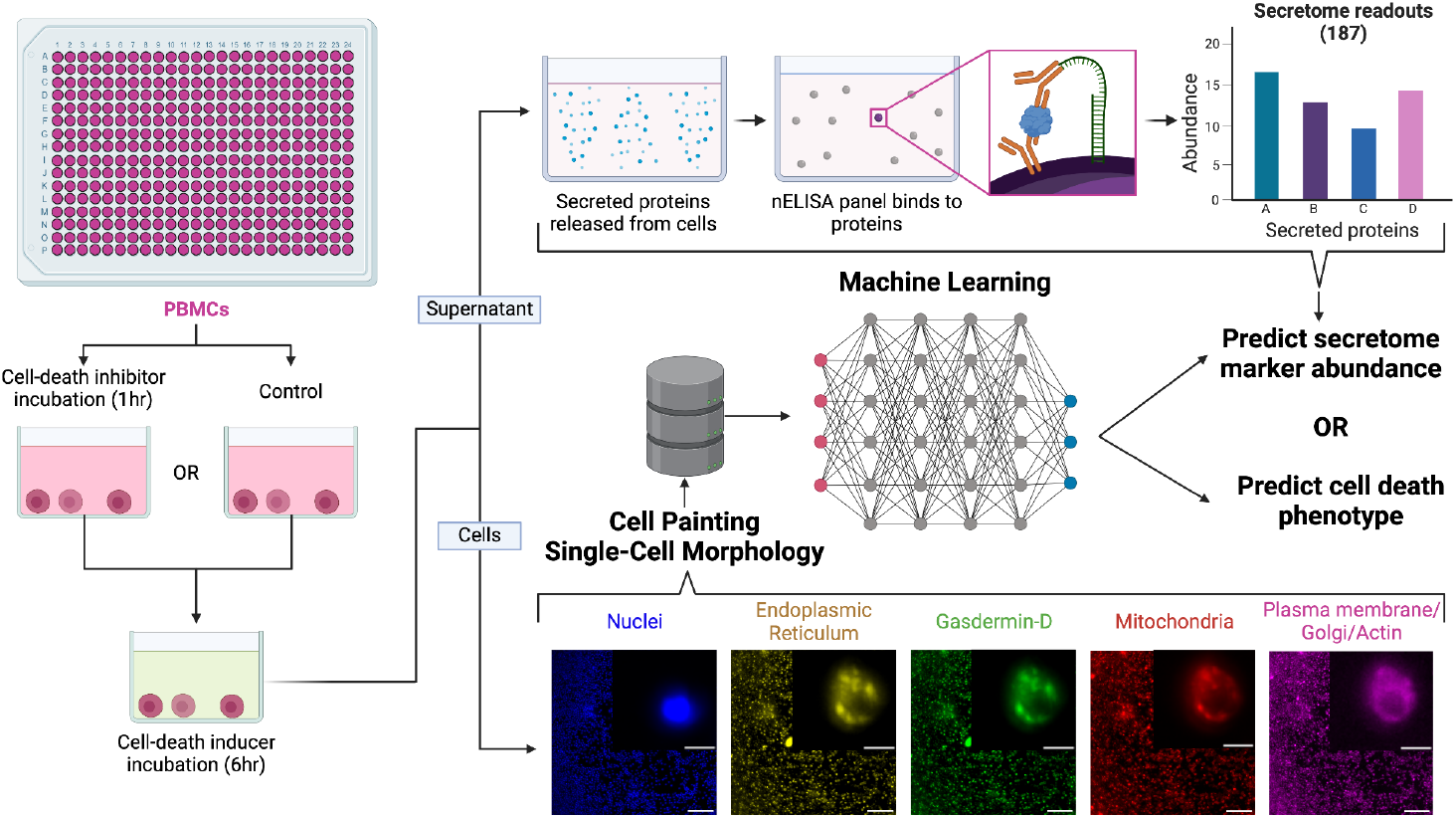
nELISA and imaging workflows. Experimental workflow for human Peripheral Blood Mononuclear Cells (PBMCs) treated with cell death inhibitors and inducers. We use the supernatant for the nELISA panel to identify the abundance of 187 secreted proteins and the fixed cells for a modified Cell Painting assay, which stains for six organelles and cleaved Gasdermin D. We applied different machine learning pipelines to predict individual secretome marker abundance and single-cell cell death phenotype. The scale bars in the representative Cell Painting image represent 5μm (inset) and 100μm (full image).

We confirmed that pyroptotic treatments were eliciting an inflammatory pyroptotic response via the secretion of canonical marker IL-1β and lacking secretion of apoptotic marker CCL24 (**Fig. 2A; Supplemental Figure 3A**).^40,56–60^ We also confirmed that apoptotic treatments did not elicit an inflammatory secretome response by measuring the TNF-α and IL-1β secretion **(Supplemental Figure 3B)**. Interestingly, by secretome markers, H2O2 treated cells clustered with the DMSO treated cells. Thus we assigned the H2O2 treated into the control group. We posit that these cells did not have enough time to respond to the H2O2 treatment thus we believe these cells to not be undergoing apoptosis. We applied Uniform Manifold Approximation Projection (UMAP)^61^ which showed that replicate wells grouped together indicating high reproducibility. The UMAP also showed apoptosis treatments clustering separately from pyroptosis treatments, which formed two clusters: One cluster driven by LPS and flagellin and the other by LPS+nigericin **(Fig. 3B; Supplemental Figure 3C**). This could be explained by pathway activation differences: LPS activates TLR4, Flagellin activates TLR5, and nigericin elicits potassium efflux. Furthermore, we observed an expected enrichment of other canonical cell death markers for specific treatments. For example, pyroptosis inducers selectively increased levels of IL-18 and IL-6, while apoptosis inducers selectively increased CCL24, CCL13, and IL-2 (**Fig. 3C**).^62–74^

**Figure 2.**
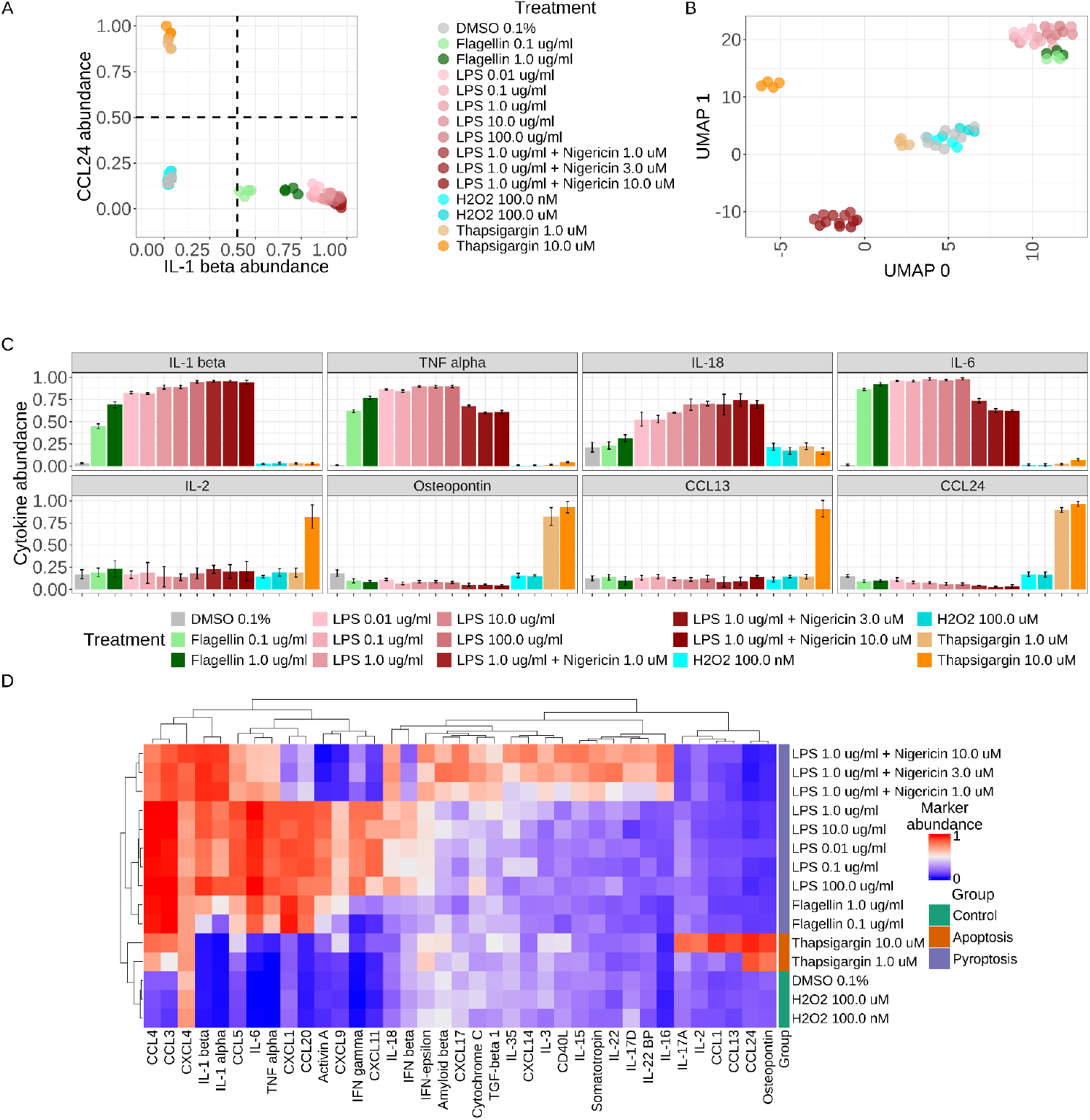
Pyroptosis and apoptosis have distinct secretome profiles for a subset of treatments. **(A)** Abundance of IL-1β and CCL24 secretions, which are canonical pyroptosis and apoptosis secretome markers. Axis values are min-max scaled abundance measurements from the nELISA assay. The dotted lines represent gates used as ground truth for treatments that induce or do not induce pyroptosis. See Supplementary Figure 3 and methods for complete cell death categorizations using canonical secreted markers. **(B)** Applying Uniform Manifold Approximation Projection (UMAP) to secretome data reveals differential clustering by specific cell death-inducing agents and controls. **(C)** Min-max normalized abundance of secreted proteins known to be associated with pyroptosis and apoptosis. Whiskers on bars represent standard deviation across well replicates. **(D)** Select secreted secretome marker values (analysis of variance [ANOVA] per feature; selected features with p < 0.05) across treatments show differential distinct secretome markers abundance for apoptotic, pyroptotic, and control treatment cells.

**Figure 3.**
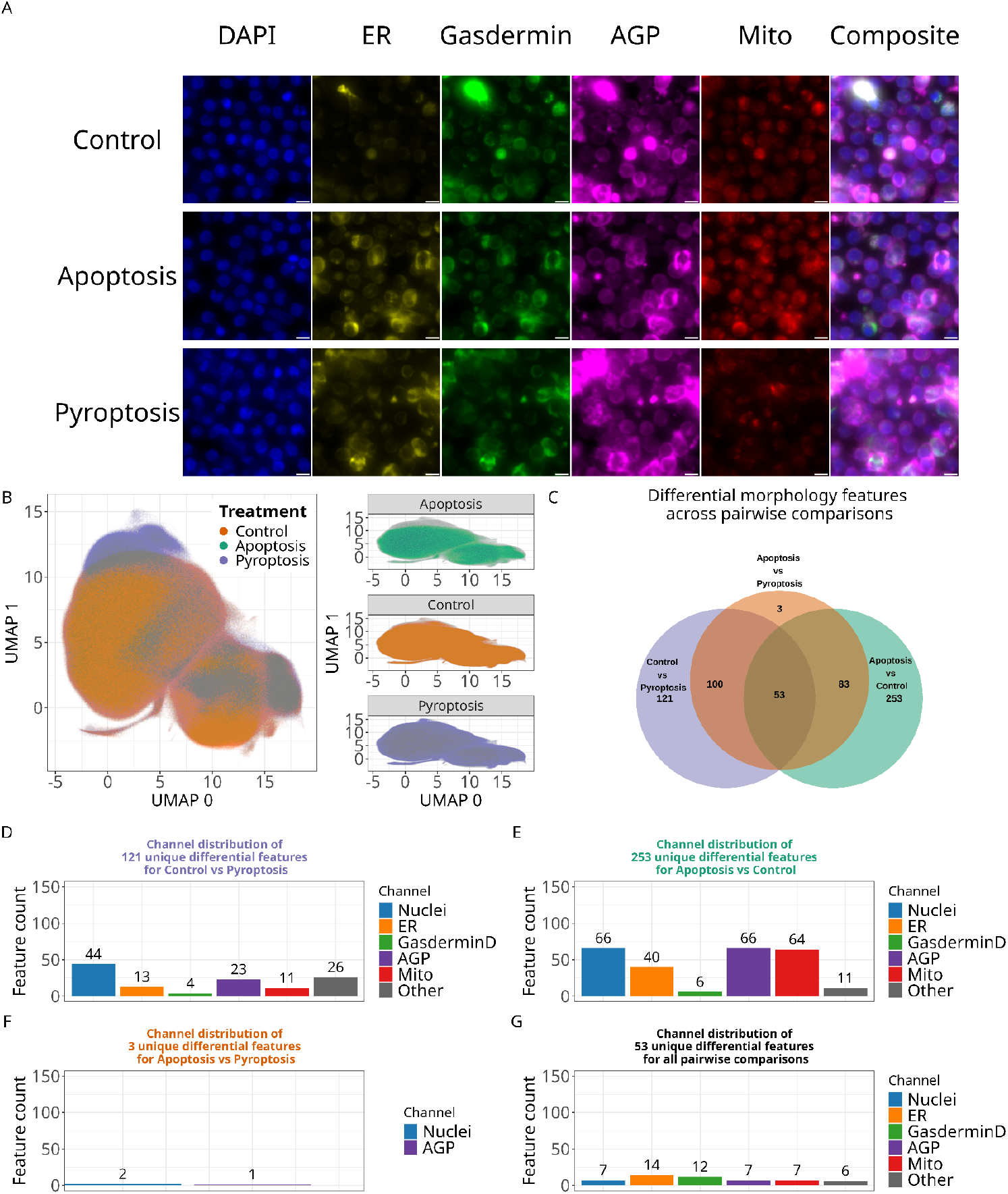
Cell Painting distinguishes single-cell morphology differences across different cell death types. **(A)** Human PBMC montage of Cell Painting. The scale bar represents 5μm. Each row represents single-cells randomly selected to represent each cell death class. Control (top row), apoptosis (middle row), and pyroptosis (bottom row). The composite images are the merged blue, green, and magenta channels, representing the DAPI, Gasdermin D, and AGP channels, respectively. **(B)** UMAP visualization of single-cell morphologies across selected treatments. **(C)** Venn Diagram of differential morphology features identified by ANOVA and Tukey’s HSD post hoc test. Tukey’s HSD identified morphology differences across channels for **(D)** differential features distinct for the pyroptosis vs. control pairwise comparison, **(E)** apoptosis vs. control, and **(F)** apoptosis vs. pyroptosis treatments. **(G)** All 420 morphology features that were different in any comparison.

We applied an analysis of variance (ANOVA) to the 187 secretrome markers independently and observed 37 differentially secreted markers between cells treated with apoptosis inducers and cells treated with pyroptosis inducers after correcting for multiple tests (Tukey’s Honestly Significant Difference [HSD] test p < 0.001). Hierarchical clustering of these secreted markers showed clear and consistent differences for each treatment group **(Fig. 2D; Supplemental Figure 4)**. Many of these differential secretome markers have not been fully reported in the literature as apoptosis-specific, including CCL24, Osteopontin, and IL-2. See **Supplementary Table 2** for a full list of significantly different secretome markers across control, apoptotic, and pyroptotic cells.

### Cell Painting reveals distinct single-cell morphologies across pyroptotic and apoptotic treatments

We also performed high-content microscopy in the same wells where we measured the secretome profiles. Specifically, we applied a modified Cell Painting assay to include a stain for N-terminus cleaved Gasdermin D (GSDMD) (**Fig. 3A**). We used CellProfiler^75^ as part of an image analysis pipeline for quality control (QC), illumination correction, cell segmentation, and single-cell morphological feature extraction. We then used CytoTable^76^ and pycytominer^77^ to perform image-based profiling for single-cells (see methods for details) (**Supplemental Figure 5**). In total, we measured 2,907 morphology features in 8,318,724 single-cells (439,316 apoptosis, 3,578,372 pyroptosis, and 4,301,036 controls).

Applying UMAP to feature-selected single-cell morphology features did not reveal a clear separation between pyroptotic, apoptotic, or control treatments (**Fig. 3B; Supplemental Figure 6**). Nevertheless, we applied an ANOVA to identify differential single-cell morphology features across pyroptotic, apoptotic, and control treatments **(Supplemental Table 2)**. Again applying Tukey’s HSD to each feature and performing multiple testing corrections across all features, we observed 613 differential features when comparing all pairwise groups, but many differential features were unique to a specific group (**Fig. 3C**). For example, we observed 121 unique differential morphology features between pyroptosis and control cells, with most morphology changes relating to the nuclei, actin, and golgi apparatus, and plasma membrane (AGP) channels (**Fig 3D**). A previous cyro-EM study identified co-localization of Golgi apparatus vesicles with cytoskeletal filaments during pyroptosis.^78^ Contrastingly, we identified 253 differential morphology features between apoptosis and control group cells, with most morphology changes from the mitochondria, nuclei, and golgi apparatus, and plasma membrane (AGP) (**Fig. 3E)**. When comparing the apoptosis with pyroptosis group cells, we identified only 3 unique differential features (**Fig 3F**). Of the 1,199 feature-selected morphology features, 53 were differential (4%) between all comparisons (**Fig 3G**). We observed a total of 613 differential features in any comparison (51%) indicating widespread morphological changes that occur during cell death. To ensure that the widespread morphological changes reported are not simply a result of the large number of cells (∼8.3 million), we performed the same ANOVA procedure we described previously using a randomly permuted feature space. This resulted in 36 (3%) differential features, indicating that our large sample size is not influencing feature difference discoveries (**Supplemental Figure 7**).

### Secretome and morphology profiling provide complementary information for mapping cell death states

We next sought to determine the complementarity of secretome and high-content cell morphology. Because the data are paired (both measurements from the same well), we can directly compare each measurement. We observed high mean average precision (mAP) scores^79^ for both morphology and secretome measurements, although secretome mAP was generally higher (**Fig 4A; Supplemental Figure 8**). mAP measures both consistency of replicate treatments and effect size compared to controls^79^, which indicates that our treatments and replicates induced large and consistent changes in secretome and morphology readouts.

**Figure 4.**
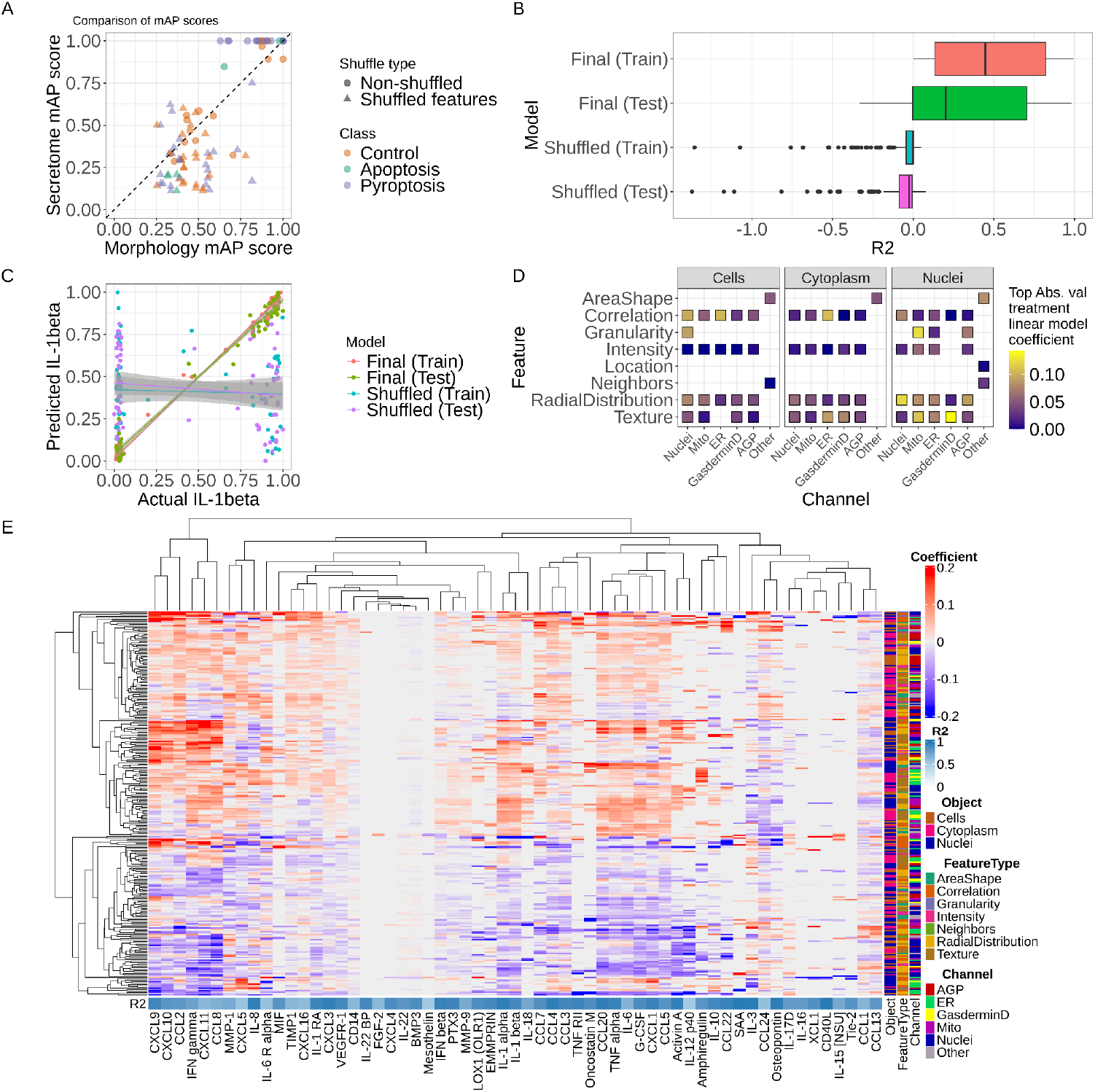
Cell morphology predicts secretome markers. **(A)** The mean average precision (mAP) of each treatment annotated by cell death type for both morphology and secretome. **(B)** The R^2^ score for every model (187) across data splits and data shuffles. **(C)** The predicted value of IL-1β compared to the actual value across train and test splits, as well as shuffled baseline and non-shuffled models (see methods), show comparatively high predictive performance. **(D)** The top absolute value logistic regression coefficient for each feature type, compartment, and channel reveals that predicting individual secretome markers uses many different kinds of morphology features. **(E)** Heatmap of the logistic regression coefficients of each selected morphology feature for each selected secretome marker model. Each model has its R^2^ score labeled, and each feature has its compartment, feature type, and channel annotated. We order the rows and columns by hierarchical clustering using complete linkage distance metric.

We next trained 187 logistic regression models with an elastic net penalty to predict individual secretome marker abundances using well-level cell morphology of all treatments. Upon testing we found that 20% of the models (40/187) had R^2^ performance in the test set greater than 0.8, indicating high performance (**Fig. 4B**). Model performance was particularly low in cases where morphology contributed to a low amount of variance in secretome marker data (**Supplemental Figure 9A**). Importantly, we see low performance in randomly shuffled data (a negative control baseline), which confirms that certain secretome markers can be predicted directly from cell morphology readouts. However, considering all secretome markers, our predictions were only marginally better than the shuffled baseline, which highlights the secretome complexity and the ability to predict only certain secreted markers from cell morphology (**Supplemental Figure 9B**). For example, IL-1β is one of the highest-performing models (test set R^2^ = 0.98) (**Fig. 4C**). CCl24, CCL4, IL-18, IL-2, IL-6, Osteopontin, and TNF-α also showed high model performance in the test set (**Supplemental Figure 9C**). Despite high predictive power globally, some predictions were particularly poor (**Supplementary Figure 9D**). For example, IL-11 test set performance was R^2^ = -0.27.

We can directly interpret the importance of individual morphology features for making secretome marker predictions. For example, accurate IL-1β predictions rely on many different morphology feature categories, particularly on granularity patterns of mitochondria near nuclei and the correlation of ER distribution with other organelles in the cytoplasm **(Fig. 4D)**. Other high performing secretome markers were influenced by diverse feature sets as well **(Fig. 4E; Supplemental Figure 10)** While the majority of secretome markers could not be predicted by morphology, most morphology features contributed to some secretome marker predictions (**Supplemental Figure 11**). The link between morphology features and secretome markers present mechanistic hypotheses about morphological changes that occur either as a cause or consequence of the processes that lead to specific marker secretion.

We next performed a Leave One Channel Out (LOCO) analysis to assess the importance of individual channels in predicting all secretome markers. We found that performance decreased, on average by 1.5%-2.8%, when we removed a channel compared to models trained with all channels (**Supplementary Figure 12A**). We observed R^2^ values between -0.36-0.99 when we removed one channel compared to R^2^ values between -0.38-0.98 for all channels (**Supplementary Figure 12B**). In addition to a negligible decrease in R^2^ values when removing a channel, we see little changes in the explained variance of models associated with their R^2^ values (**Supplementary Figure 12C**). Most importantly, when specifically removing the GSDMD channel for predicting IL-1β, we saw marginally decreased performance (**Supplemental Figure 12D**). Overall, this analysis verifies that 1) a single channel alone is not contributing to secretome predictions, 2) including all channels improves the ability of our models to predict secreted markers, and 3) cell morphology features in channels other than GSDMD can predict pyroptosis.

### Machine learning predicts cell death phenotypes in single-cells

To better understand single-cell differences in pyroptosis and apoptosis, we trained a fully-connected, two-layer Multi-Layer Perceptron (MLP) using single-cell morphology data to predict the cell death category. We defined the cell death categories at the treatment level using IL-1β and CCL24 secretion levels, as shown in **Fig. 2A**. We used training, validation, testing, and hold-out data splits (see methods) to evaluate the performance of our model on single-cells, including cells treated with agents that the model had never before seen (**Supplemental Figure 13**). Our models show high precision and recall, indicating generalizable predictions with minimal overfitting, specifically for predicting pyroptotic and control cells (**Fig. 5A**). We see a much lower predictive performance for apoptosis, likely because of having far fewer single-cells and treatments. Confusion matrices and F1 scores further show particularly high performance for pyroptosis predictions, even in held-out wells (**Fig. 5B**; **Supplemental Figure 14)**. Single-cell class probabilities are skewed toward the correct cell death mechanism, representative single-cell images show differential morphology features that align with canonical understanding by eye (e.g., Gasdermin D translocation to membranes) (**Fig. 5C-E; Supplemental Figure 15**). Importantly, the entire treatment hold-out set (flagellin 0.1 ug/mL, flagellin 1ug/mL, flagellin 1ug/mL + 1uM disulfiram) performed with an F1 score above 0.75 compared to its shuffled baseline F1 score of less than 0.15 **(Supplemental Figure 16A)**. We show the predicted probability distribution for single-cells treated with the flagellin hold outs and a predicted probability of one representative image in **Supplemental Figure 16B-C**.

**Figure 5.**
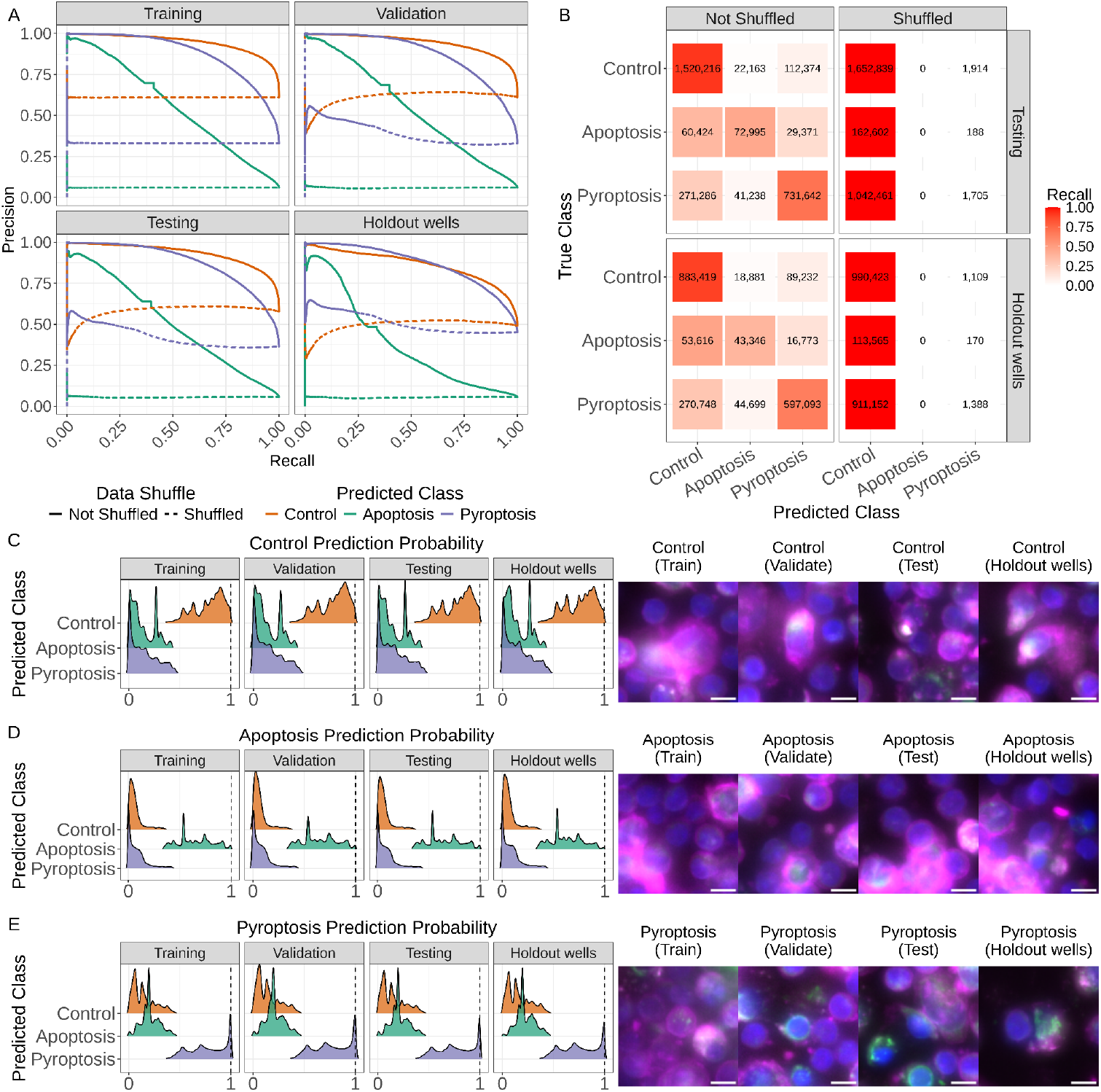
Machine Learning predicts single-cell death states. **(A)** Precision-recall curves of the training, validation, testing, and hold-out well data splits for both the non-shuffled and shuffled baseline models. Confusion matrices for shuffled and non-shuffled models showing the testing and hold-out well data splits. The number in each square represents the number of predicted single-cells, and the color represents the recall of each class. Probabilities for **(C)** control, **(D)** apoptosis, and **(E)** pyroptosis predictions stratified by true class. A single-cell cropped image for cells with a probability equal to 1 for each class and data split; where the channels represent: AGP = magenta, Gasdermin D = green, and DNA = blue. AGP is the actin, golgi apparatus, and plasma membrane stain in the canonical Cell Painting panel. The Gasdermin D is the N-terminal cleaved region. Scale bars represent 5μm.

## Discussion

In treatment-stimulated human PBMCs, paired secretome and high-content imaging demonstrated a complementary map that characterizes pyroptosis. The secretome data confirmed canonical pyroptotic markers (IL-1β and TNF-α), but it also revealed a more dynamic landscape that differs even between pyroptosis treatments. Specifically, the LPS plus nigericin-treated cells showed a different pyroptotic secretome than the flagellin or LPS-alone-treated cells. This could result from the differential activation of TLR4 (LPS),^74^ potassium efflux via ionophore channel (nigericin)^80^, and/or TLR5 activation (flagellin).^74^ We observed high levels of IL-6, IL-1 beta, IL-18, and TNF-alpha secretion in wells treated with pyroptosis inducers, which is consistent with previous studies. ^81,82^ We also unexpectedly observed that our apoptotic cells secreted the inflammatory markers IL-2, CCL13, CCL1, CCL24, and osteopontin, which contradicts previous descriptions of the non-inflammatory nature of apoptosis.^83,84^ However, other recent research categorizes these markers as anti-inflammatory and anti-apoptotic^66,72,85,86^ or even inflammatory in certain circumstances such as caspase inhibition and mitochondria-initiated apoptosis.^7^ Both apoptosis and pyroptosis share similar secreted markers, implying that many markers are indicative of general cell stress and cell death, or if they are also present in the control group, that they are part of normal human PBMC processes (see **Supplemental Figure 4**). Perplexingly we also reported that H2O2 treated wells clustered with the DMSO treated wells. We cannot explain why this occurred, but we still included the H2O2 treated wells in all analyses in our control group. We verified that these cells were most likely not undergoing necroptosis due to levels of TNF-alpha almost identical to that of DMSO treated wells.^87,88^

Our high-content imaging and image-based profiling approach characterized the cell morphology landscape of pyroptosis. Pyroptosis altered nuclei and AGP morphology, while apoptosis had more morphology feature changes, mostly impacting nuclei, AGP, and mitochondria. Deep neural networks trained using single-cell morphology features robustly predicted cell death class, even in never-before-seen wells and treatments. Recent approaches have used deep learning to predict cells dying by apoptosis and ferroptosis.^37^ The paired secretome data combined with our modified Cell Painting panel to include Gasdermin D, added important layers of ground truth to the imaging data, which enabled us to explore important links between data types. For example, we observed that machine learning models can accurately predict many individual secreted markers using diverse morphology features from different organelles. This suggests a mechanistic link between form and function, but the causal directionality remains unknown (i.e., do single-cell morphology changes indicate preparation for secretome marker secretion, or does secretome marker secretion cause cells to change morphology?). Overall, our map provides a multi-modal, systems biology characterization of pyroptosis, which reveals a complex secretome and morphological landscape that overlaps and diverges from apoptosis.^37^ While we were revising this work, two other studies, which applied similar machine learning approaches to identify targets of pyroptosis, were presented.^25,89^ These new results corroborate our results, confirming the ability to perform image-based profiling on cells undergoing pyroptosis. Given that pyroptosis is extensively less studied compared to apoptosis **(Supplemental Figure 17)**, our map of pyroptosis provides two major advances for drug discovery and cell death biology: a) enabling further mechanistic studies to build on organelle-defined image-based profiles, and b) objective elucidation of key differences between apoptosis and pyroptosis.

There are many limitations to our approach. First, we collected data only six hours after treatment induction, which provided only a snapshot of the early stages of pyroptosis. A cell experiencing pyroptosis might commit to cell death or upregulate proliferative pathways that rescue cells from death.^90,91^ Nevertheless, a six hour incubation point distinctly separates pyroptosis from apoptosis in both the secretome and cell morphology. We aim to further elucidate the intracellular dynamics of cells undergoing forms of cell death, such as pyroptosis and apoptosis, in a temporal-dependent manner in future studies. Second, we also measured only 187 secretome markers and thus missed the full secretome response. However, with technical advances, 187 markers are more secretome markers than previously imaginable via ELISA-based assays, and we expect content to increase in the future. Cell secretion can be variable and pleiotropic, with multiple secreted secretome markers being indicative of multiple biological processes. Thus we were cautious of oversimplified interpretations due to the multifaceted nature of cell secretion profiles. Third, our secretome marker profiling only measures total markers secreted by all single cells within a well. Therefore, we are not able to distinguish which cells are secreting which markers, and it is possible that certain cells are secreting different combinations or a disproportionate quantity of secretome markers. However, despite this potential heterogeneity, we still see high performance in using well-aggregated cell morphology to predict many individual secretome markers. Fourth, although a well might show evidence of a pyroptotic inflammatory response, every cell might not necessarily be undergoing pyroptosis. This could result in false negative classifications of single cells in our deep learning analysis since ground truth is assigned at the well level. Fifth, the Cell Painting assay used five imaging channels, and including the Gasdermin D stain introduced some spectral overlap. We combat this issue via feature selection methods that remove highly correlated features across channels (see methods). Nevertheless, our machine learning and comparative analyses identify differential features across all imaging channels, indicating that while by eye they may appear similar, our computational methodology captures different features. Sixth, we do not control for cell heterogeneity as we lack ground truth knowledge of the ratio of cell types contained within our human PBMC population. Our machine learning approaches take this heterogeneity into account, and nevertheless perform robustly, even in never-before-seen populations. And finally, seventh, we are using a relatively small dataset. We treated human PBMCs from a single donor with 36 treatment combinations in 154 individual wells collected in a single 384-well plate. While this yielded approximately 8.3 million segmented single cells, our analyses that aggregated information at the well level reduced our dataset to 154 well samples. Despite this reduction, we were still able to predict certain secretome markers from morphology in never-before-seen cells and treatments, which reliably revealed important morphology indicators that were different across cell death classes. It is likely that donor-to-donor differences exist and different cell types could reveal different pyroptosis maps. Furthermore, there is likely plate-to-plate variation that might also contribute to differences, and we expect future studies will expand and refine this pyroptosis map by also exploring additional cell lines, treatments, and adding temporal content.

## Conclusion

We applied a systems biology approach to better understand pyroptosis, a less studied type of PCD, which allowed us to characterize cell morphology changes and relate them with specific inflammatory secretome markers. Increasing knowledge about pyroptosis can have profound impacts in the pathophysiology of different diseases and the potential to accelerate drug development.

## Materials and Methods

### Cell culture

We cultured human PBMCs in a controlled environment of 37ºC with 5% CO2. We plated human PBMCs in a consistent medium at an approximate density of 125,000 cells per well on Poly-L-Lysine coated plates to encourage adherence (see **Supplemental File 1** for specific conditions). We incubated human PBMCs for 30 minutes prior to compound treatment. We also applied a similar cell culture, data collection, and experimental strategy to SH-SY5Y cells, although we do not report these results in this manuscript.

### Perturbation panel

We treated cells with 36 different agent combinations dissolved in DMSO (**Supplemental Table 1**). We incubated cells for one hour with cell death inhibitors/controls, then incubated these cells for six hours in cell death inducers/controls. We included six to eight replicate wells of each inhibition/induction combination (36 combinations) within one 384-well plate. For a complete plate map, see **Supplemental Figure 2**. We chose this panel to induce canonical and non-canonical pyroptosis, generic cell death, and canonical apoptosis.

### Optimizing time points to capture dying but not dead cells

We cultured PBMCs and treated them with DMSO 0.1%, LPS 1.0 μg/mL, LPS 10 μg/mL, Thapsigargin 1 μM, and Thapsigargin 10 μM. We incubated cells with each perturbation for three, six, and nine hours. We added propidium iodide at a final concentration of 0.1 mg/mL, and Hoescht at a final concentration of 2 μM. We determined the number of dead cells using propidium iodide staining, while counting the total number of cells using Hoechst. For each condition, we imaged five fields of view using an epifluorescence microscope at 10X magnification. We calculated the number of live cells by subtracting the number of dead cells from the total number of cells. To determine the proportion of live cells, we divided the number of live cells by the total number of cells. We calculated the percentage of dead and live cells for each perturbation at each specified time point.

### Cell Painting

We modified the canonical Cell Painting assay to remove the cytoplasmic RNA and nucleoli stain for a stain that specifically marks cleaved N-terminal Gasdermin D (Anti-cleaved N-terminal GSDMD Rabbit Monoclonal, Abcam).^28^ Additional Cell Painting organelles and stains included Nuclei (Hoechst 33342); the endoplasmic reticulum (Concanavalin A PhenoVue Fluor 488); Golgi apparatus and the plasma membrane (WGA PhenoVue Fluor 555); actin (Phalloidin PhenoVue Fluor 568); and mitochondria (PhenoVue 641 Mitochondria stain).

We followed the version 3 Cell Painting protocol.^92^ Briefly, at six hours post compound incubation, we treated the cells with live cell mitochondria stain for 30 min at 37°C, 5% CO2. We then fixed cells in 30 μL of 5.33% (w/v) at a final concentration of 4% (w/v) of methanol-free PFA in PBS for 20 minutes at room temperature. We then permeabilized cells in a permeabilization buffer for 5 minutes. Following permeabilization, we added 50 uL of blocking solution for one hour at room temperature. We stained cells with Gasdermin D primary antibody overnight at 8°C. We then stained cells with secondary antibody and Hoechst for one hour at room temperature. We incubated cells in permeabilization buffer and Cell Painting dyes for 30 minutes at room temperature. We subsequently imaged cells with all channels **(Supplemental File 1)**.

### Image acquisition

We collected microscopy images using the Revvity (previously PerkinElmer) Operetta CLS imaging platform with a 20x water immersion NA 1.0 objective with two separate acquisitions. Our first acquisition was prior to the addition of Cell Painting stains. We acquired two channels (Hoechst 33342 and Gasdermin D) at 16 different fields of view (FOV) per well over tgree z-planes spaced out 2μm apart. After adding Cell Painting stains, we acquired images over five channels at 16 different fields of view (FOV) per well at four different z-slices spaced out 2μm apart. Channel excitation and emission wavelengths can be found in **Supplemental Table 4**. We binned pixels at 1×1. We output all Cell Painting in 16-bit TIFF format, preserving data integrity through lossless compression.

### Image analysis

We used CellProfiler^75^ for image analysis. We applied maximum projection, which takes the highest pixel intensity value per pixel position from all four z-slices. Once there was one z-stack per FOV, we corrected the images for illumination errors, such as vignetting, using the standard CellProfiler approach.^93^ We created one illumination correction function per channel and saved these functions as NPY files. We applied these functions to images during the feature extraction pipeline.

Using CellProfiler, we segmented three compartments (nuclei, cells, and cytoplasm) for every single cell. CellProfiler creates binary masks for each compartment per FOV, which are then applied to each of the five channels. We extracted features from each compartment, such as granularity, texture, intensity, and more. We stored feature data in a SQLite database file for downstream image-based profiling. See https://github.com/WayScience/pyroptosis_signature_image_profiling for access to our CellProfiler pipelines.

### Image-based profiling

We formatted the extracted features from CellProfiler by performing image-based profiling via the Cytomining software ecosystem **(Supplemental Figure 5)**. We used software called CytoTable^76^ to process and clean the CellProfiler SQLite output file. CytoTable merges SQLite compartment tables to create one row per single-cell and outputs the data into the high-performance Apache Parquet format.^94^ We then used pycytominer software to perform the rest of the image-based profiling pipeline.^77^ The pipeline consists of first annotating the single-cells with metadata from a plate map file associated with each single-cell (e.g., treatment, dose, etc.). We then normalized the features using the per-cell type negative controls (DMSO-treated cells) as a reference using the standardized method (z-score normalization). We performed feature selection on the normalized data to remove redundant or non-informative features. Feature selection consisted of removing features that are highly correlated with other features (Pearson correlation coefficient of 0.9 or higher), removing blocklist features known to be noisy^95^, removing features with low variance (features that had greater than 10% of samples with the same measurement), and removing features that had more than 5% of samples with missing values. In summary, we segmented and measured 8.3 million single cells and extracted a total of 2,907 features per single-cell (we retained 1,199 features after feature selection).

### nELISA panel

Following the Cell Painting data acquisition, we isolated the supernatant from the same treatment wells, which we sent to Nomic Bio (Montreal, Canada) for nELISA-based secretome marker analysis, as described previously.^38^ Briefly, the nELISA pre-assembles antibody pairs on spectrally encoded microparticles, resulting in spatial separation between non-cognate antibodies, preventing the rise of reagent-driven cross-reactivity, and enabling multiplexing of hundreds of ELISAs in parallel. The nELISA assay measured 187 secretome markers (see **Supplemental Table 3**), resulting in paired Cell Painting and secretome profiling data at the well level. We processed the nELISA signals using min-max scaling.

### Ground truth gating via canonical cytokines

We applied a double gate against IL-1β (0.5) and TNF-α (0.5) to identify wells that included pyroptotic cell death.^40,56–60^ We deemed double positive wells to be pyroptotic and double negative wells to be non-pyroptotic. Further, we applied a double gate for IL-1β (0.4) and CCL24 (0.5) to assign apoptotic cell death.^40,86^ We considered CCL24 positive and IL-1β negative wells as apoptotic. We considered all wells that were negative for both gates as controls. These processes aligned with our cell death expectations given the existing literature about all compounds.

### ElasticNet regression models framework

We trained and evaluated ElasticNet logistic regression models using sklearn v1.3.0.^96,97^ We used mean-aggregated well morphology data to predict 187 secretome markers from 1,199 feature-selected morphology features as inputs (normalized data before feature selection). We trained individual models per secretome marker. Further, we trained shuffled baseline models to use as negative controls. To avoid overfitting, we split our data stratified by treatment. We split 25% of the lowest and highest LPS doses (0.010 ug/mL and 100 ug/mL) to the training set and 75% to the testing set. We kept LPS at 1 ug/mL, both with and without nigericin at 3uM, 100% in the testing set. This provides a never-before-seen dose to use in the testing set for assessing model generalizability. In addition, we kept all doses of flagellin in the testing set. For all remaining treatments, we performed a 50% training and 50% testing split. We trained the models using a leave-one-out cross-validation (LOOCV) approach and optimized the following model parameters from the following search spaces. We optimized α parameters [0.001, 0.01, 0.1, 1, 10, 100, 1000, 10000] and the L1/L2 ratio [0.01, 0.1, 0.2, 0.3, 0.4, 0.5, 0.6, 0.7, 0.9, 0.99] where L1 = 0 and L2 = 1. We also trained and tested on a permuted feature space to establish a shuffled baseline.

### Leave One Channel Out (LOCO) ElasticNet regression models

The LOCO analysis follows the same methods described above in the ElasticNet regression models framework section, with the exception of one channel being removed during both training and testing. It is important to note that we removed all features associated with a specific stain, which included correlation features that related to other stains. Leaving in these correlation features poses a higher risk for information leakage from left out channels to influence performance. We performed LOCO systematically for every channel, which resulted in five times as many additional elastic net models to evaluate.

### Multi-layer perceptron training framework

We designed and trained a multi-class, two-layer, Multi-Layer Perceptron (MLP) to predict cell death types.^98^ We used the Pytorch v2.0.0 library for the model creation, training, and testing.^99^ We used Optuna v3.3.0^100^ to optimize model hyperparameters from the search space described in **Supplemental Table 5**, which also provides the optimized architecture and hyperparameters. We also trained and tested on a permuted feature space to establish a shuffled baseline. To rigorously evaluate performance, we used a data splitting procedure that resulted in three different holdout sets, each with increasing generalizability difficulty. First, we held out all single-cells from one well for every treatment we used for training. This would tell us the generalizability of never-before-seen wells. Second, we held out all single-cells from the following treatment doses: LPS 1ug/mL and LPS 1ug/mL + Nigericin 3uM. This would tell us the generalizability of never-before-seen doses of a treatment the model did see. Lastly, we held out all single cells treated with flagellin (Flagellin 0.1 ug/mL, Flagellin 1ug/mL, Flagellin 1ug/mL + 1uM Disulfiram). This would tell us the generalizability of a never-before-seen treatment.

We split cells that received the 0.01 ug/mL and LPS 100 ug/mL treatments into 75% and 25% testing and training sets respectively. We split the remaining single-cells by 50% into testing and training data splits. We further split the training dataset into a 20% validation set, which we used to determine early-stopping criteria during training and thus avoiding overfitting. We balanced our training data split by treatment to ensure data splits represent distributions of the original data. We split the LPS treatments, which contained many more single cells, differently from the other treatments to retain a balance of treatments in the training set. Single-cell counts by treatment and data split can found in **Supplemental Table 6**. We also evaluated performance on randomly permuted features to compare as a negative control.

### Representative image selection

We selected a representative image using a predefined metric for all image montages. In this study, we used the predicted probability of a cell’s predicted cell death class as the metric of choice to correctly represent predicted cells from our machine learning model (see **Fig. 5**). We used images of cells that had a predicted probability of 1.0.

### Morphology and secretome ANOVA

We determined differential morphology features and secreted markers using a series of ANOVAs. We grouped cells treated with controls, apoptotic inducers, and pyroptotic inducers as defined in **Fig. 2A**. We applied an ANOVA for each feature individually. For the morphology analysis, we also included the number of single cells per well as a covariate to adjust for morphology differences that are known to occur with differing cell densities. We further performed a Tukey’s Honestly Significant Difference [HSD] test to adjust p-values for multiple testing and determine specific group differences.^101^ We considered any feature with an adjusted p value of 0.001 as significant for morphology features and adjusted p values of 0.05 as significant for the secretome features. We also randomly permuted features and separately ran ANOVAs as described above.This random permutation tells us how many features this procedure is expected to identify by chance.

### UMAP visualization

We used UMAP to visualize the dimension reduction of morphology and secretome features.^61^ We generated all UMAP plots with six nearest neighbors, a minimum distance of 0.8, and 2 components. We used the cosine similarity metric with a spread of 1.1 and random initialization.

### Mean Average Precision (mAP) analysis

Mean Average Precision (mAP) is a metric of how similar a given group is to replicates within the group in comparison to a reference group.^79^ In our case, we used mAP in multiple ways, creating different groups for comparison. Specifically, we used mAP to evaluate 1) the consistency of perturbations within the same cell death categories, 2) the consistency of individual treatments with each cell death category, and 3) the consistency of individual treatment replicates for both secretome and morphology profiles. We also applied mAP to data after applying randomly permuting features. In every case, we used DMSO treatment replicates as the reference group. We calculated mAP using the Copairs Python package.^79^

### Computational resources

We utilized the Alpine high-performance computing resource at the University of Colorado Boulder. Alpine has 382 compute nodes with 22,180 cores, 36 NVIDIA a100 GPUs, and 24 AMD MI100 GPUs. In addition to Alpine, all other computation was performed on a local workstation with the following specifications: AMD Ryzen 9 5900x CPU - 12 cores, with 128 GB of DDR4 RAM, and GeForce RTX 3090 TI GPU.

## Supporting information

Supplemental File 1 - Cell Culture Conditions

## Acknowledgments

This work utilized the Alpine high-performance computing resource at the University of Colorado Boulder. Alpine is jointly funded by the University of Colorado Boulder, the University of Colorado Anschutz, Colorado State University, and the National Science Foundation (award 2201538). We would like to thank Cameron Mattson, Roshan Kern, and Erik Serrano for performing code review. We would also like to thank Haruhiko Bito, Takanari Inoue, Francisco Quintana, Consuelo Walss-Bass, and Venetia Zachariou for advice at various stages of this project.

## Data and Code Availability

All data and code are publicly available. The raw images, illumination correction files, and extracted feature profiles were deposited to the Image Data Resource (IDR) (https://idr.openmicroscopy.org) under accession number idr0160.

All image analysis pipelines for segmentation, feature extraction, and image-based profiling are available at https://github.com/WayScience/pyroptosis_signature_image_profiling.^102^

All data analysis code is available at: https://github.com/WayScience/pyroptosis_signature_data_analysis.^103^

