## Supplemental Figures 1-17 for "A morphology and secretome map of pyroptosis"

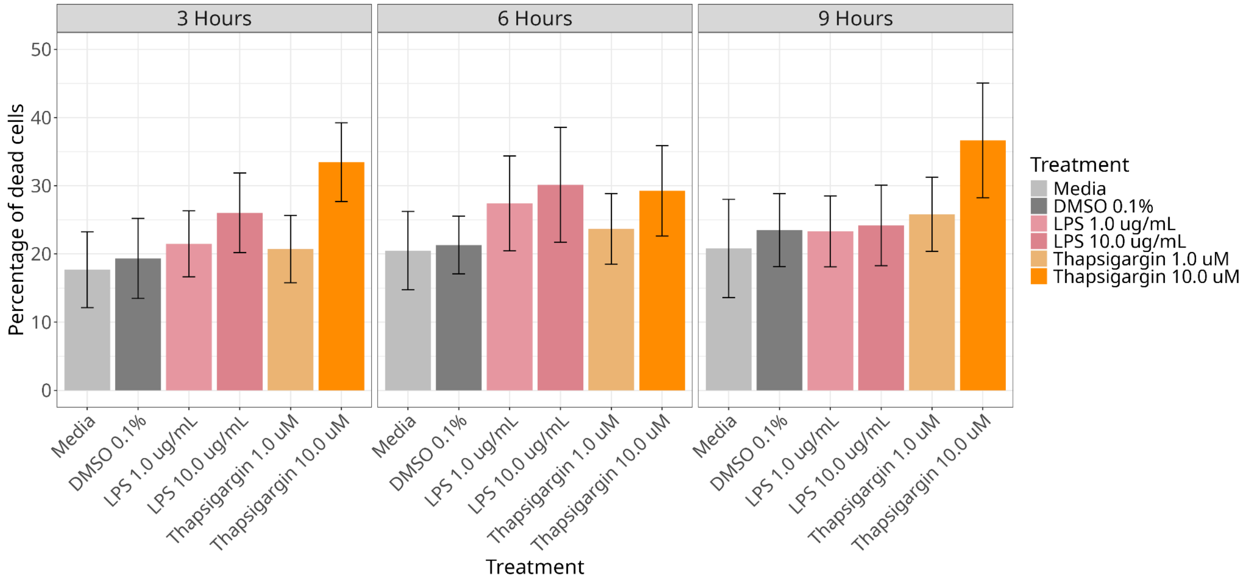

**Supplemental Figure 1:** Percentage of dead cells *in 3, 6, and 9 hour incubation with pyroptosis and apoptosis inducers.*

Lipopolysaccharide (LPS) and thapsigargin induced cell death at a higher rate compared to DMSO and media control conditions. The six hour time point revealed that while many cells were dead (~30%), the majority of cells treated with these conditions were alive or in the process of dying. These death curves are independent data from the Cell Painting data generated.

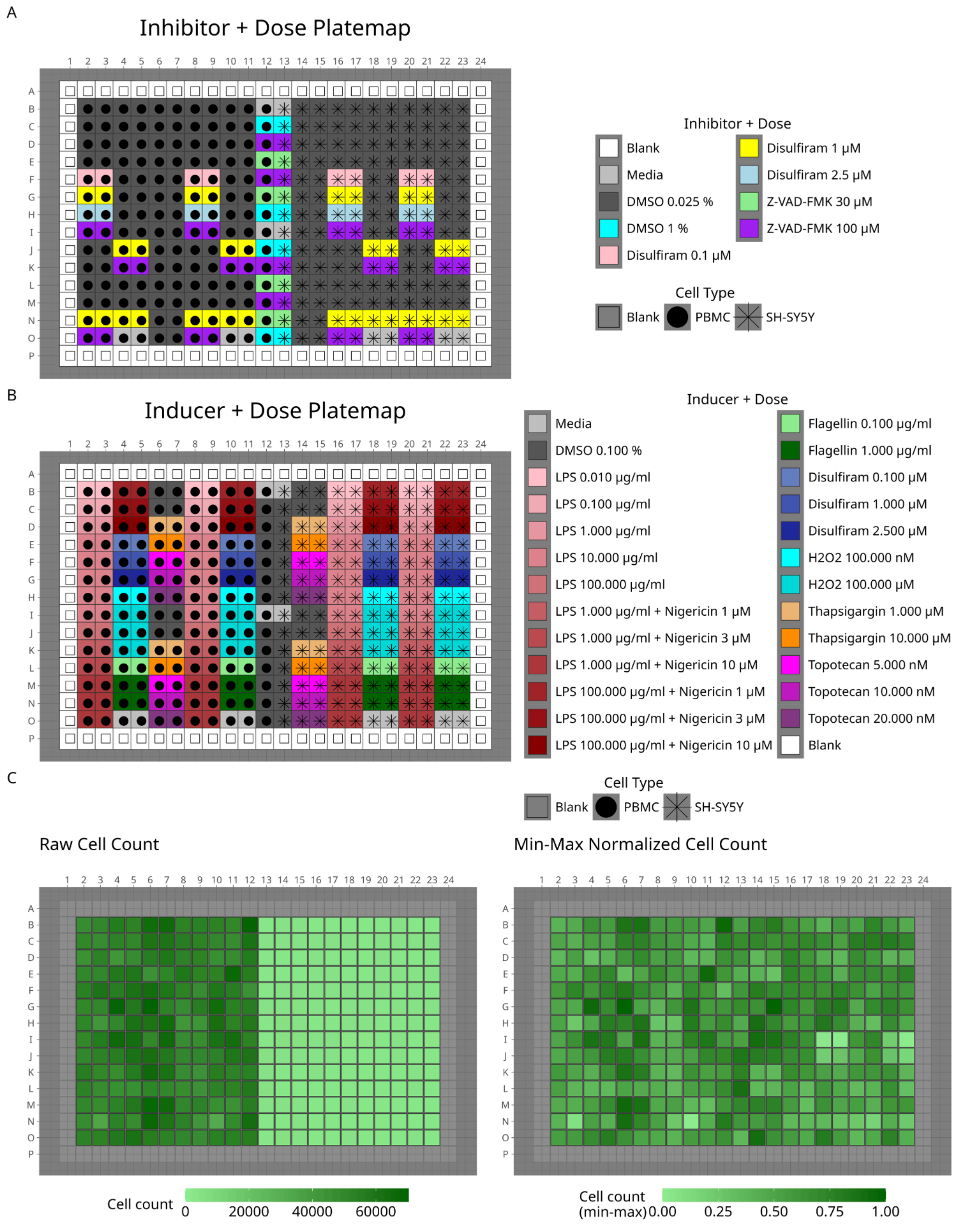

**Supplemental Figure 2:** *Plate maps of the inducers and inhibitors.*

We collected one 384-well plate of data. Half of the plate was peripheral blood mononuclear cells (PBMCs), and the other half was SH-SY5Y (neuroblastoma). We only report the PBMC data from this experiment in this paper. **(A)** We applied specific cell death inhibitors one hour prior to inducers. **(B)** We incubated inducers for six hours prior to applying the Cell Painting assay. **(C)** We also include the raw cell counts and the normalized cell counts for each cell type thus displaying variation in cell count for the different treated wells.

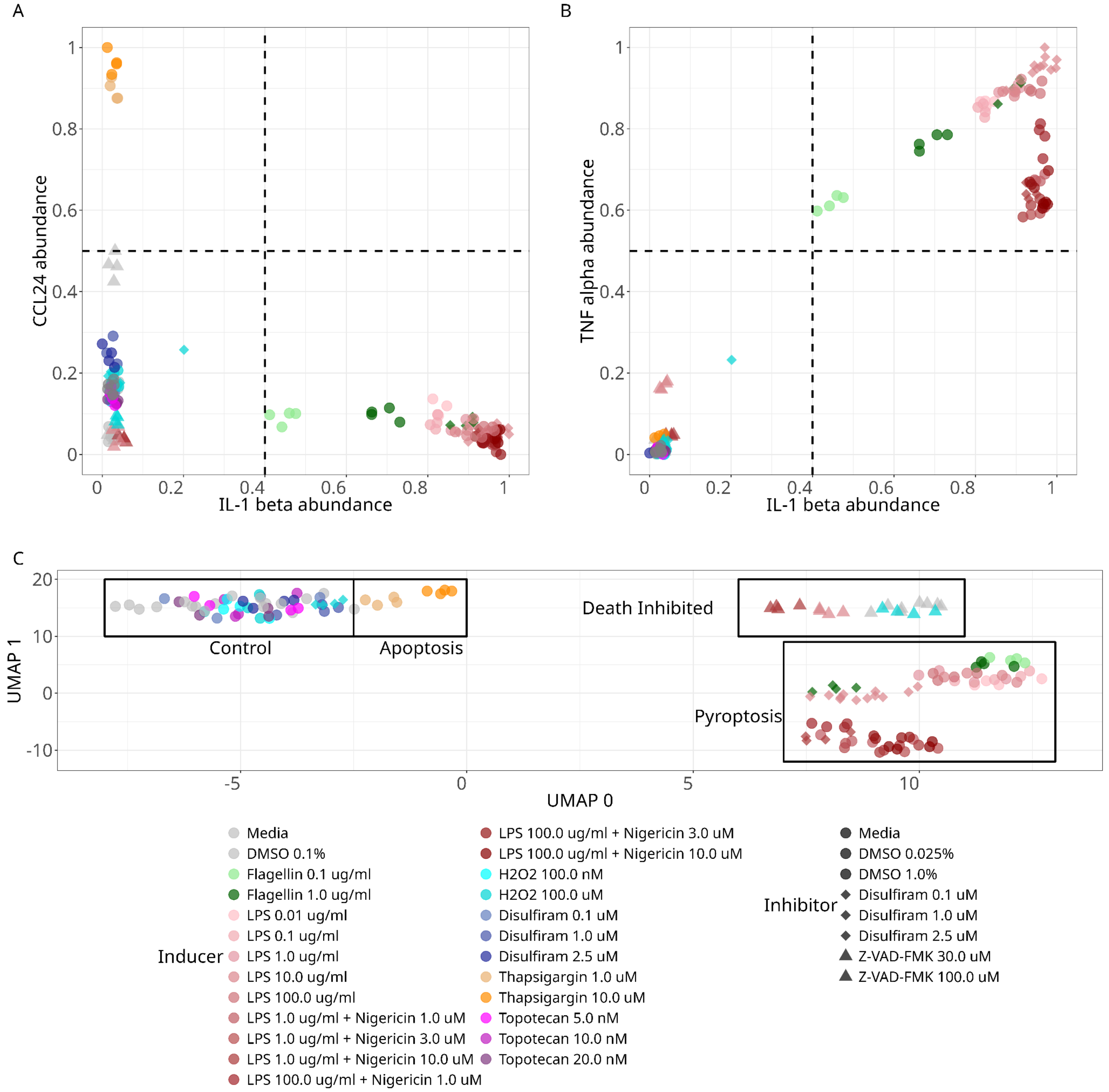

**Supplemental Figure 3:** *Confirming expected secretome grouping across all treatment combinations.*

**(A)** CCL24 - IL-1β cytokine double gate to classify pyroptotic (lower right) and apoptotic (upper left) treatments applied to all treatment combinations. **(B)** TNFα - IL-1β double gate applied to all treatment combinations. The upper right quadrant indicates treatments that induce pyroptosis. The dotted lines (for both panels A and B) indicate an x-intercept of 0.4 and a y-intercept of 0.5, which we used as gates to distinguish apoptosis from pyroptosis. **(C)** Uniform manifold approximation (UMAP) of all 187 secretome markers across all treatments.

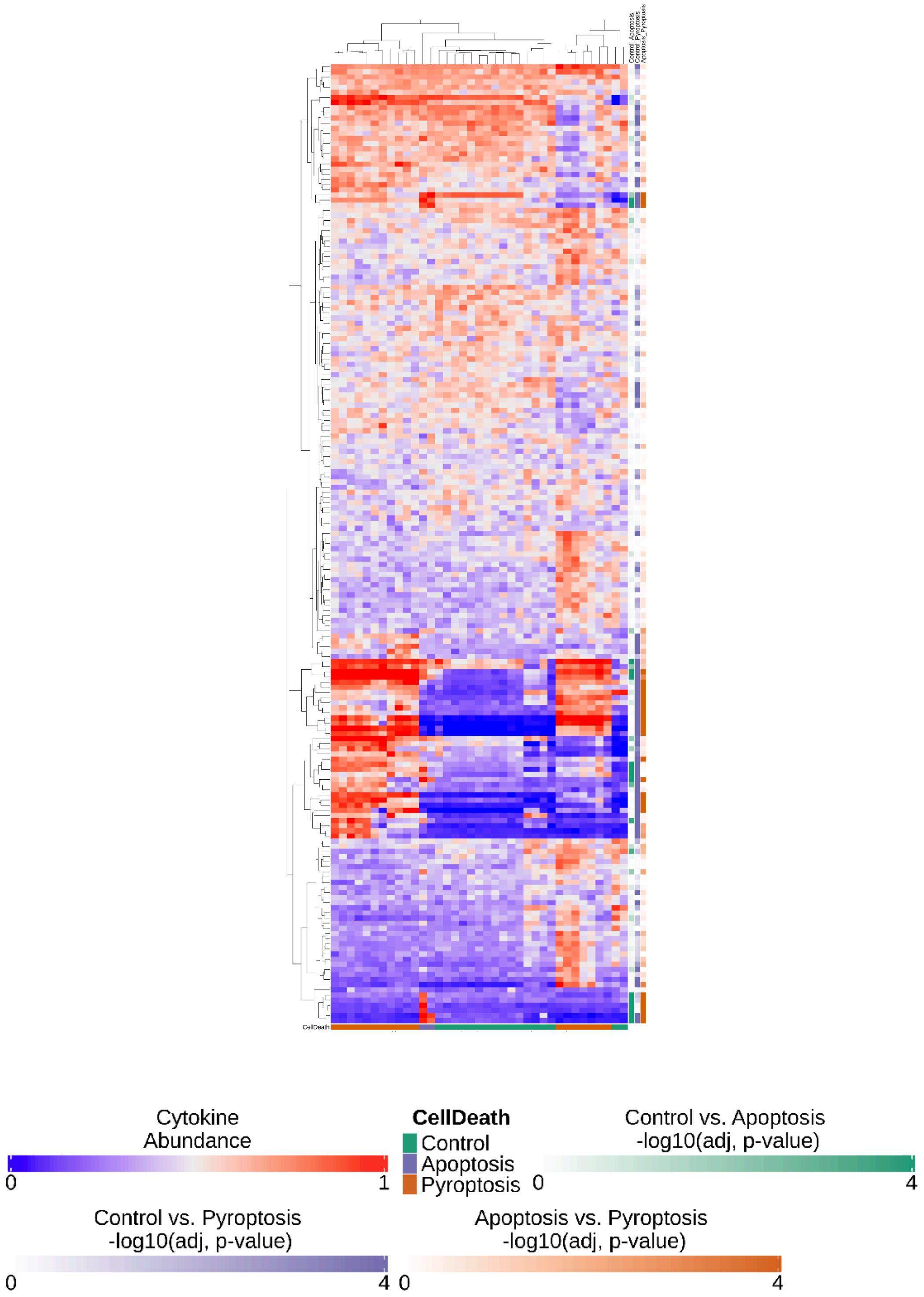

**Supplemental Figure 4:** *The complete 187 secretome profile for all treatment combinations.*

Rows are hierarchically clustered with complete linkage distance metric. Each row is annotated with the negative log10(p-value) of a Tukey's HSD test across the three cell death classes for all secretome markers. Columns represent each treatment.

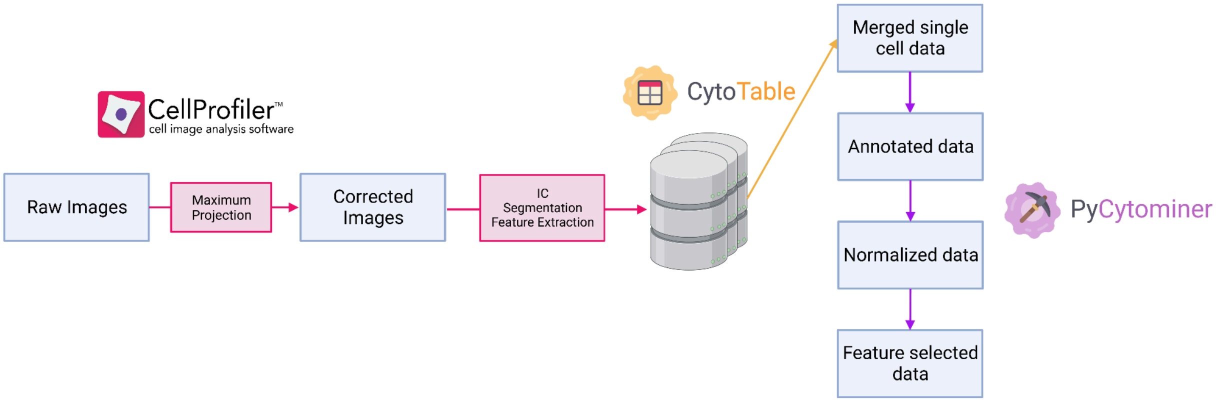

**Supplemental Figure 5:** *The full image analysis and image-based profiling pipeline..*

We developed an image analysis pipeline using CellProfiler to process images and extract image-based profiles. We then applied CytoTable and pycytominer to perform image-based profiling.

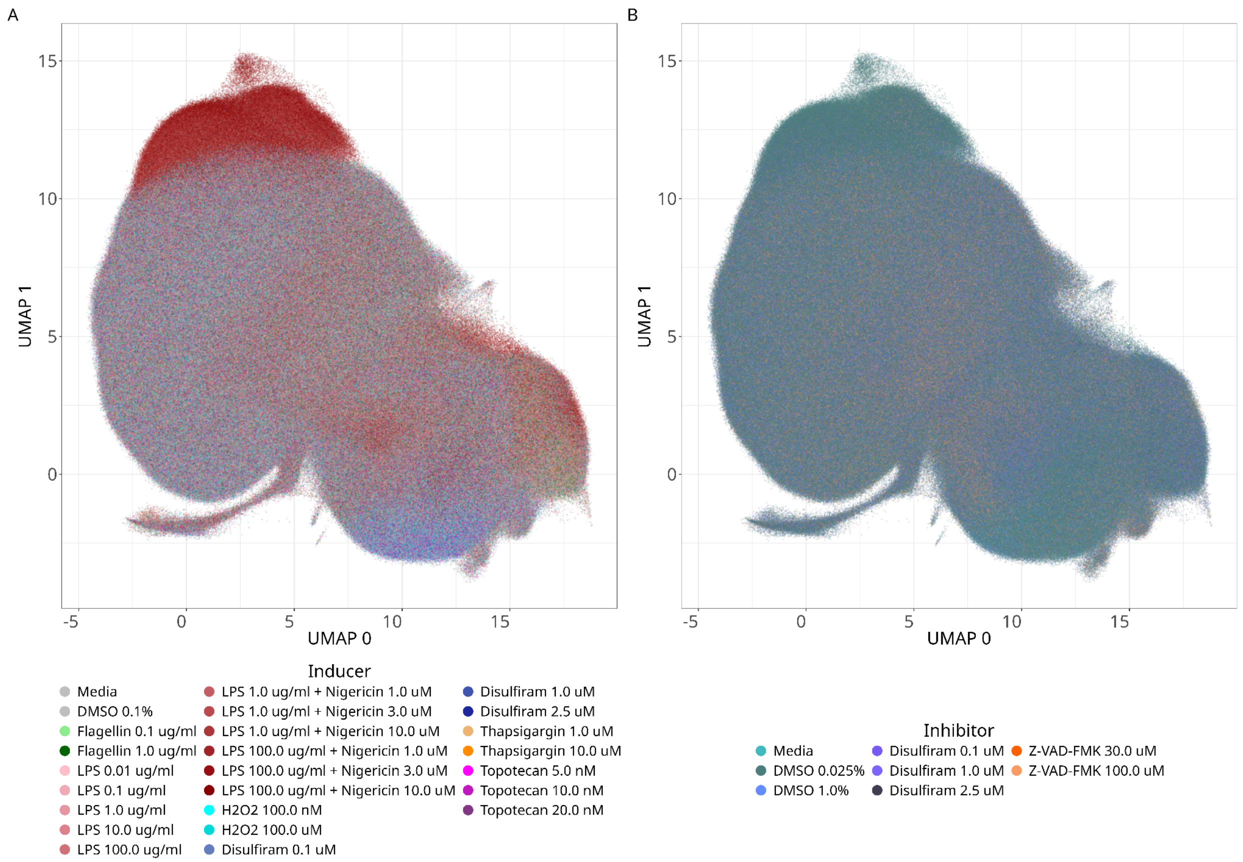

**Supplemental Figure 6:** *UMAP representations of morphology feature profiles across all treatments.*

We show the UMAP space for all inducers and inhibitor combinations used in this study. Each point represents a single-cell.

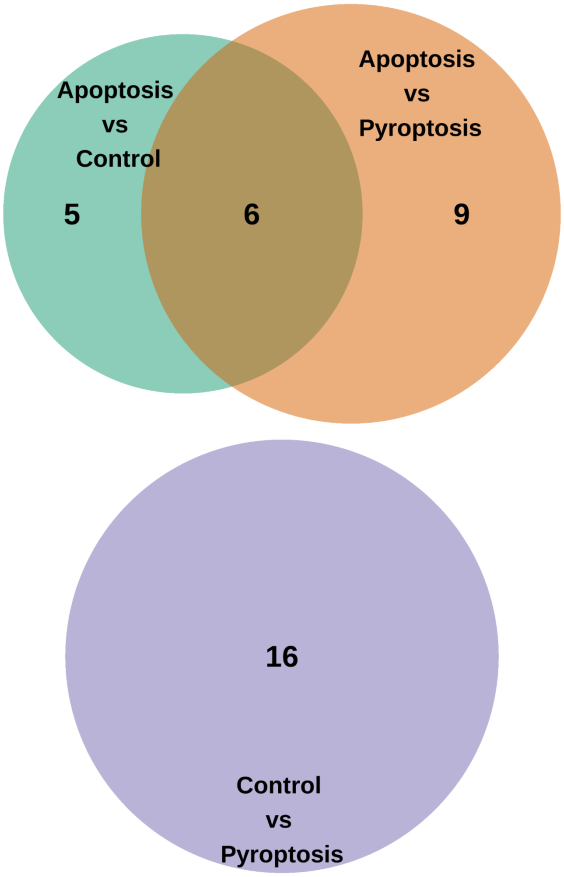

**Supplemental Figure 7:** *ANOVA of randomly permuted morphology feature space shows few differential features.*

Venn Diagram of permutation shuffled differential morphology features identified by ANOVA and Tukey's HSD post hoc test.

A

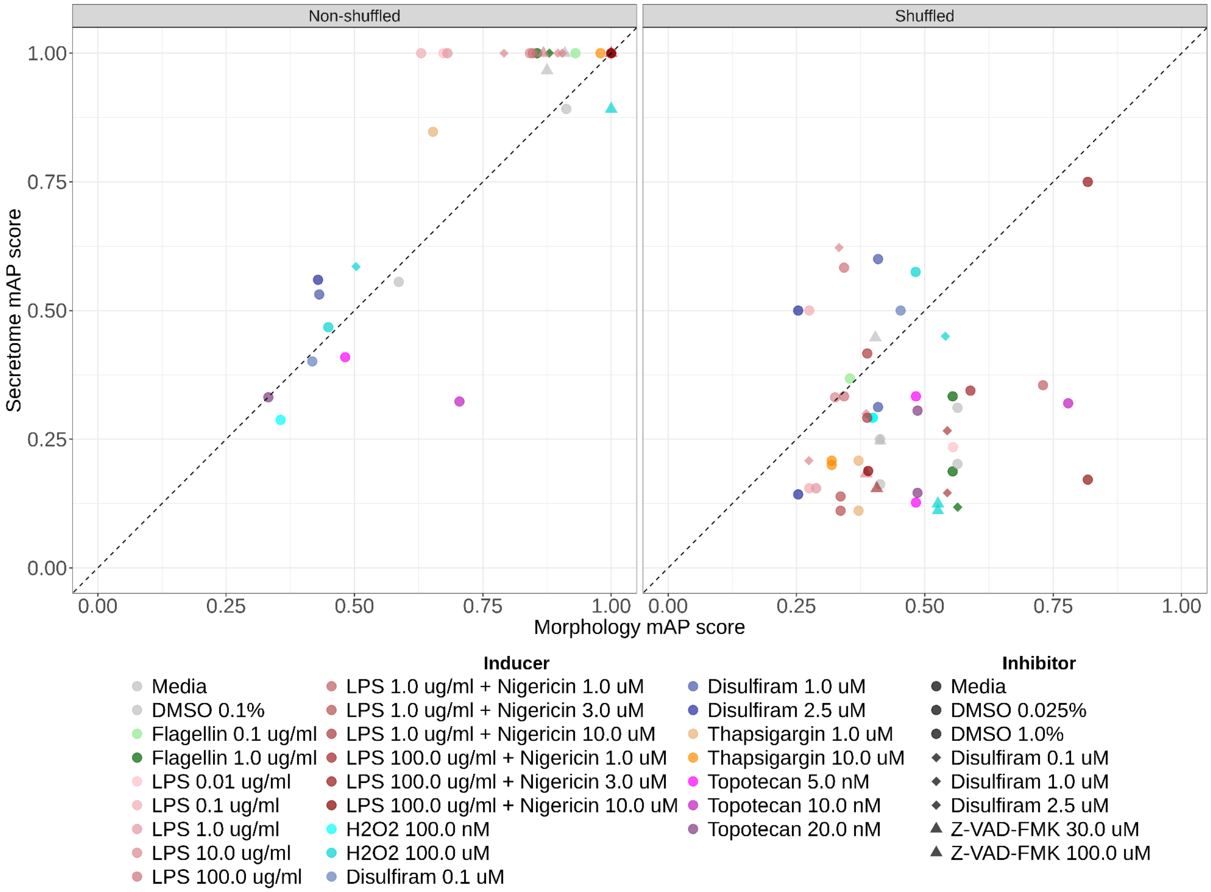

**Supplemental Figure 8:** *Mean average precision (mAP) analysis comparing secretome and morphology.*

The mean average precision (mAP) of the secretome data vs the morphology data across all treatment replicates.

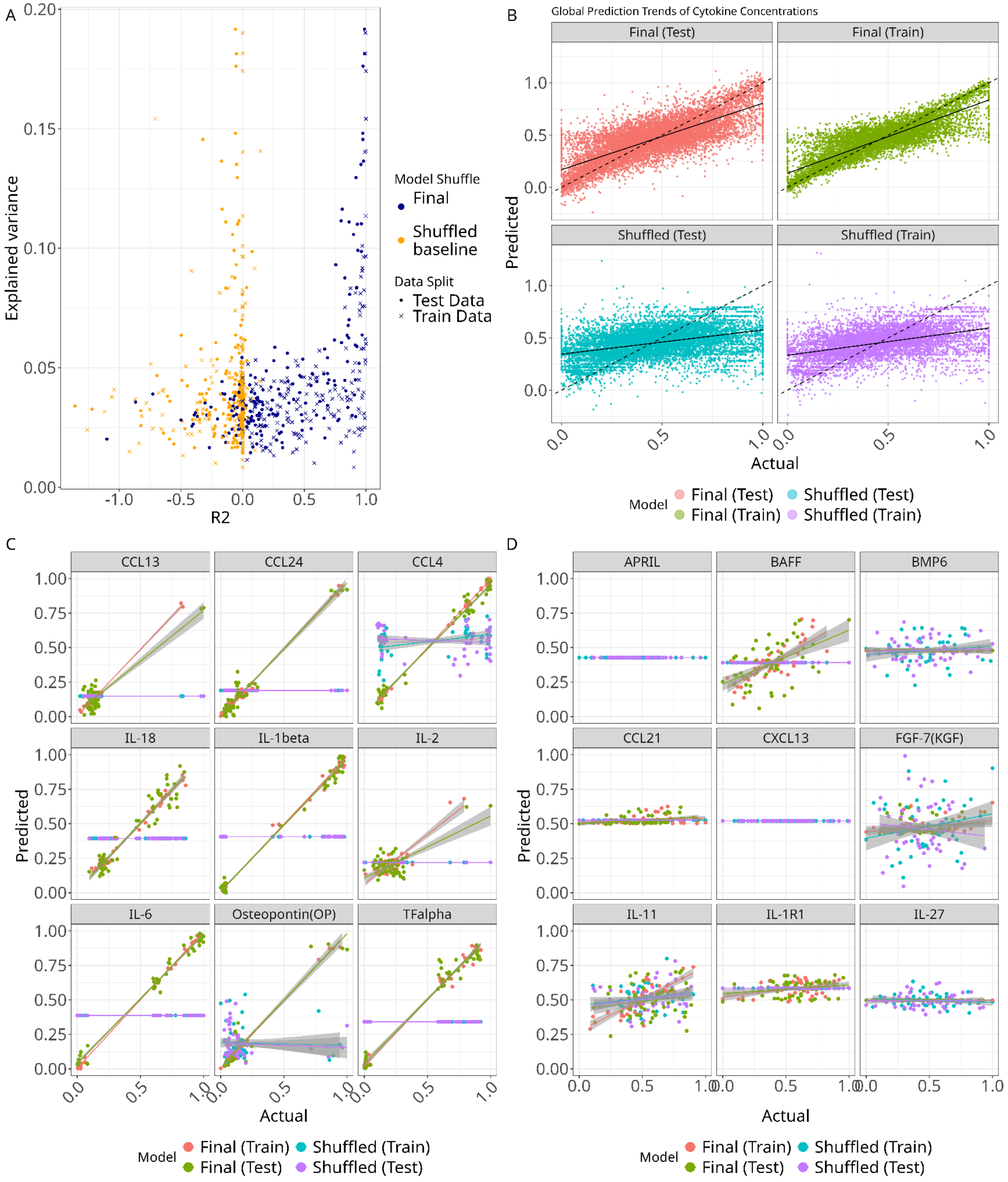

**Supplemental Figure 9:** *Machine learning model performances and expanded predictions.*

These models predict a secretome marker from the mean-well aggregated cell morphology feature space. **(A)** The explained variance against the R2 across testing and training data splits, and across shuffled and non-shuffled models. Each point represents one model. **(B)** The global (all) model predictions vs the actual value for each data split and shuffle. **(C)** The line of best fit for the global predictions against their actual levels for high-performing models. **(D)** A per-marker view of the predicted marker abundance against the actual abundance for low-performing models.

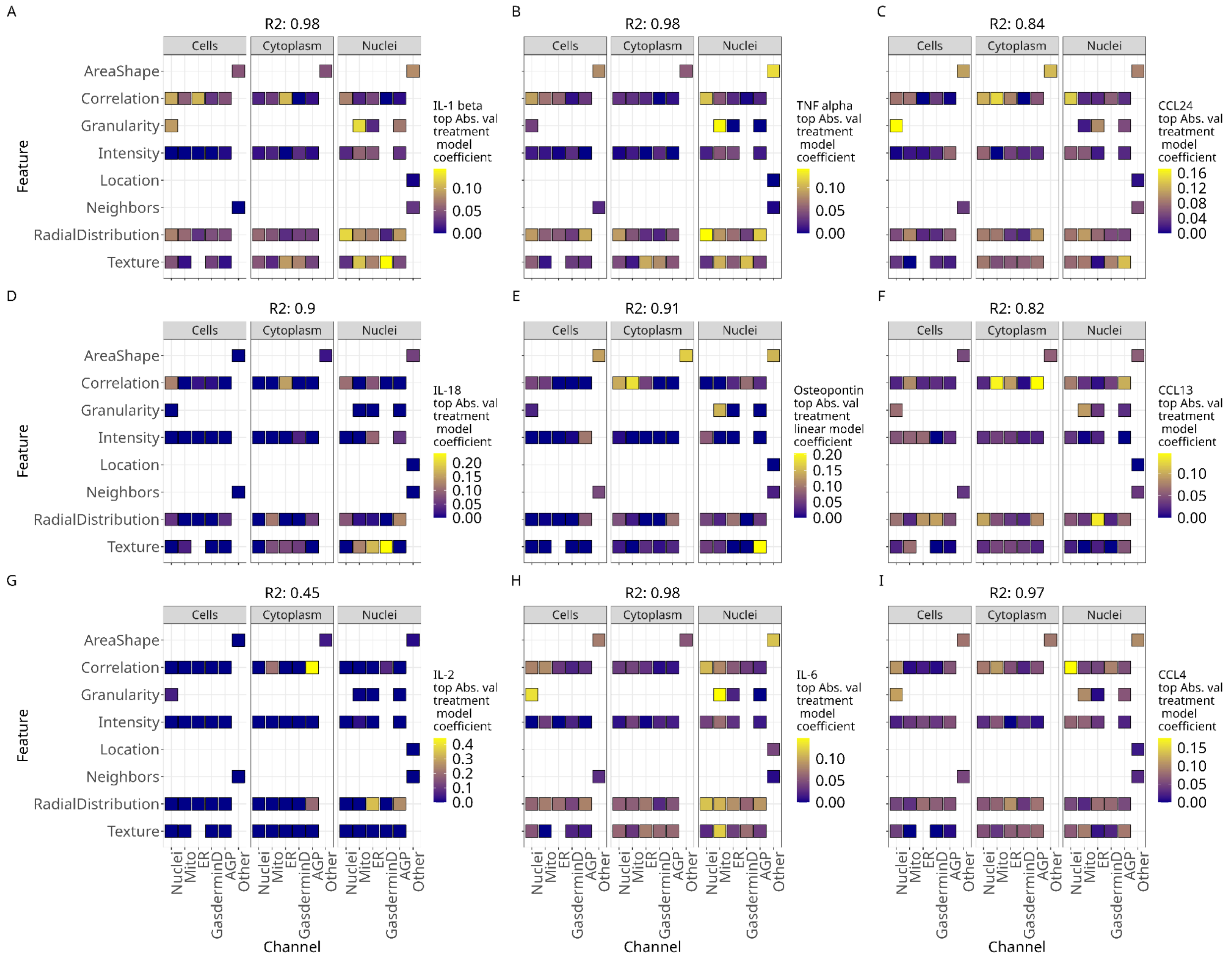

**Supplemental Figure 10:** *Model coefficients for predicting individual secretome markers with cell morphology readouts.*

Top absolute value logistic regression coefficients for **(A)** IL-1β **(B)** TNFα **(C)** CCL24 **(D)** IL-18 **(E)** Osteopontin **(F)** CCL13 **(G)** IL-2 **(H)** IL-6 and **(I)** CCL4. All models had high prediction performance (we show test set $R^2$ performance above each plot facet). We show model importance scores across cells, cytoplasm, and nuclei, imaging channels, and feature groups.

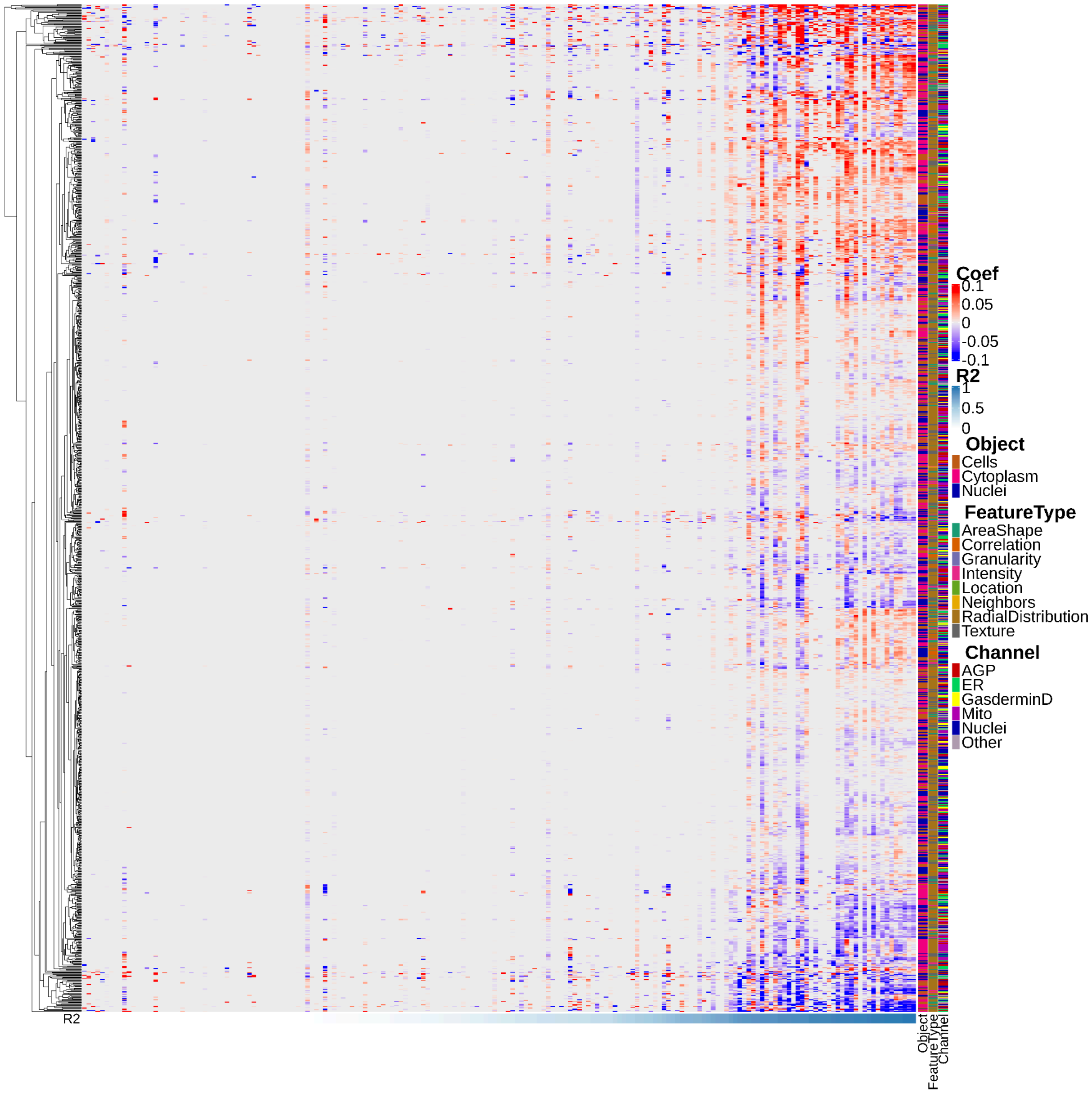

**Supplemental Figure 11:** *Morphology feature space logistic regression model for predicting individual secretome marker abundance.*

All morphology features (y-axis) against all 187 secretome markers (x-axis) represent the logistic regression model coefficients for each combination. The morphology feature type, channel, and segmentation compartment (cells, cytoplasm, or nuclei) are also visualized.

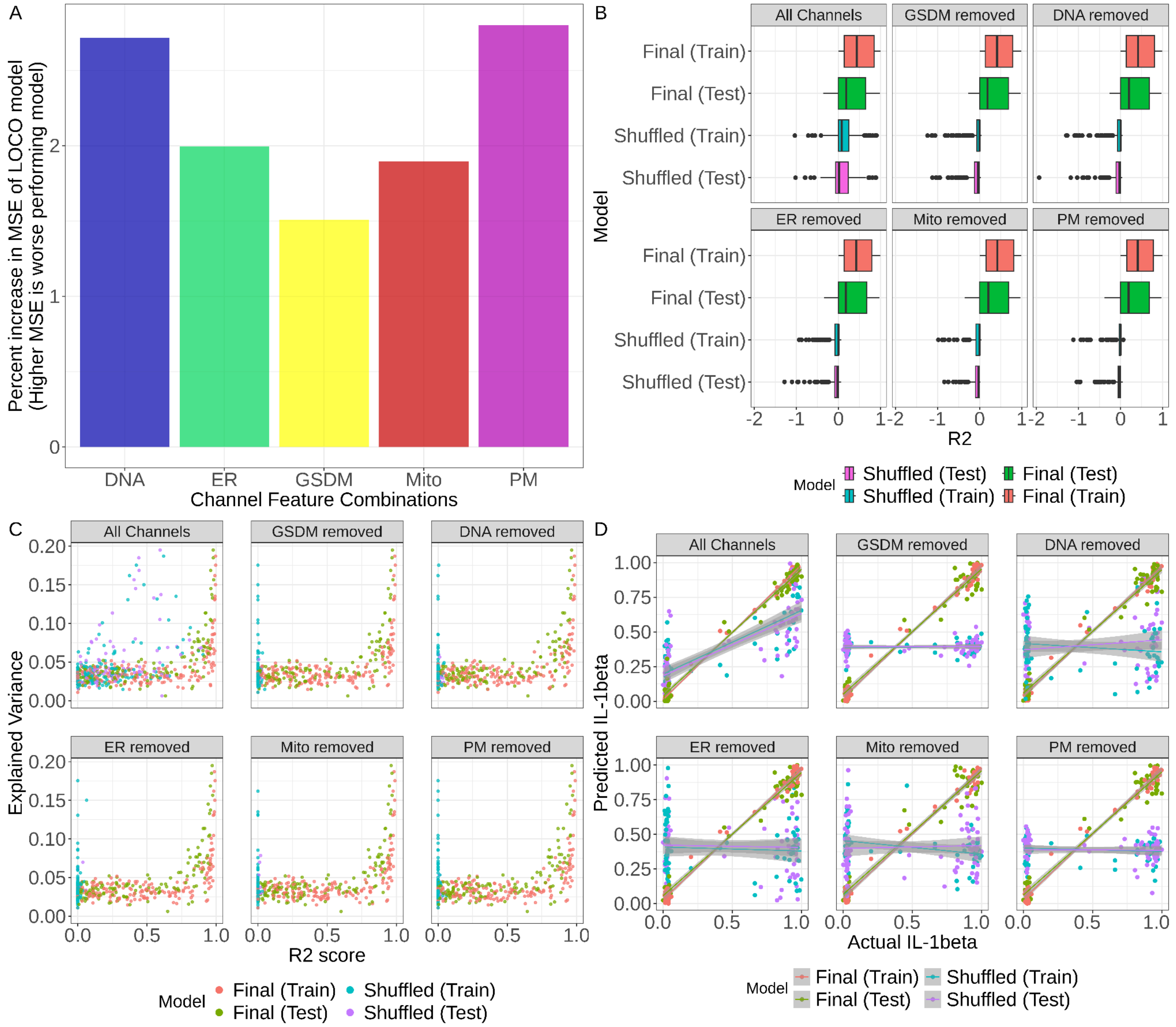

**Supplemental Figure 12:** *Leave one channel out analysis for predicting the secretome from image-based profiles.*

**(A)** The percent increase in the negative Mean Square Error (MSE) of each model for each channel that was removed for the leave one channel out (LOCO) analysis. **(B)** R2 values for all predicted secretome markers across data splits and data shuffles for each channel removed. **(C)** The explained variance against the R2 score (cut off at 0) for each data split and data shuffles for each channel were removed. **(D)** The predicted abundance vs the actual abundance of one example secretome marker, IL-1β, across all data splits and data shuffles for each channel removed.

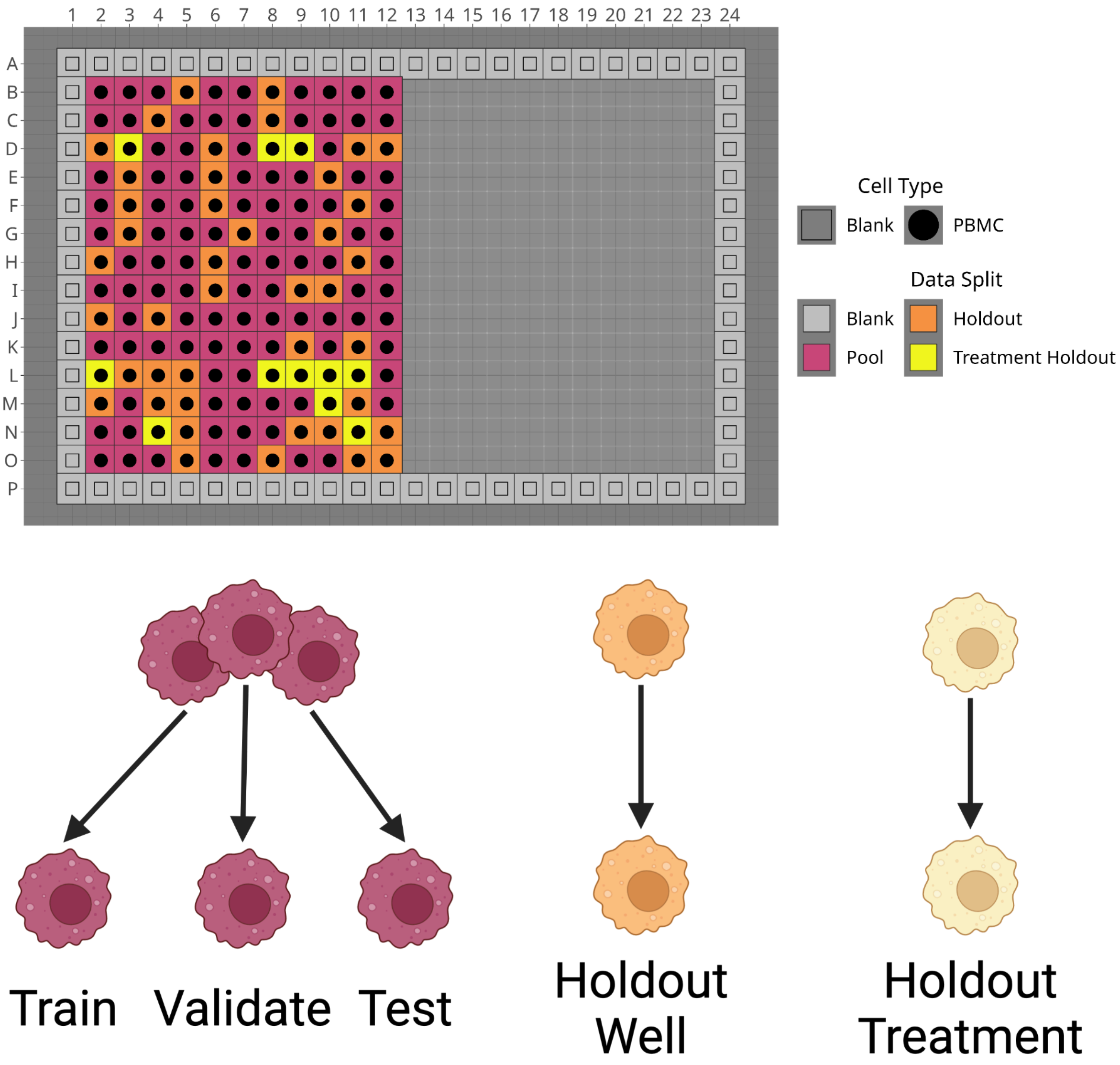

**Supplemental Figure 13:** *Our randomized data splitting procedure and plate map for single-cell multilayer perceptron.*

We split single cells in the "pooled data" wells (red) into training, validation, and testing sets. We also hold out entire wells (orange) and entire treatments (yellow) from the training procedure: (LPS at 1.0 ug/mL, LPS at 1.0 ug/mL + Nigericin at 3.0 uM, and all Flagellin treatment combinations: Flagellin at 0.10 ug/mL and 1.0 ug/mL, Flagellin at 1.0 ug/mL + Disulfiram at 1.0uM). The well holdouts are to ensure that the model is not overfitting on wells. The treatment holdout is to ensure the model is not overfitting across cell death classes.

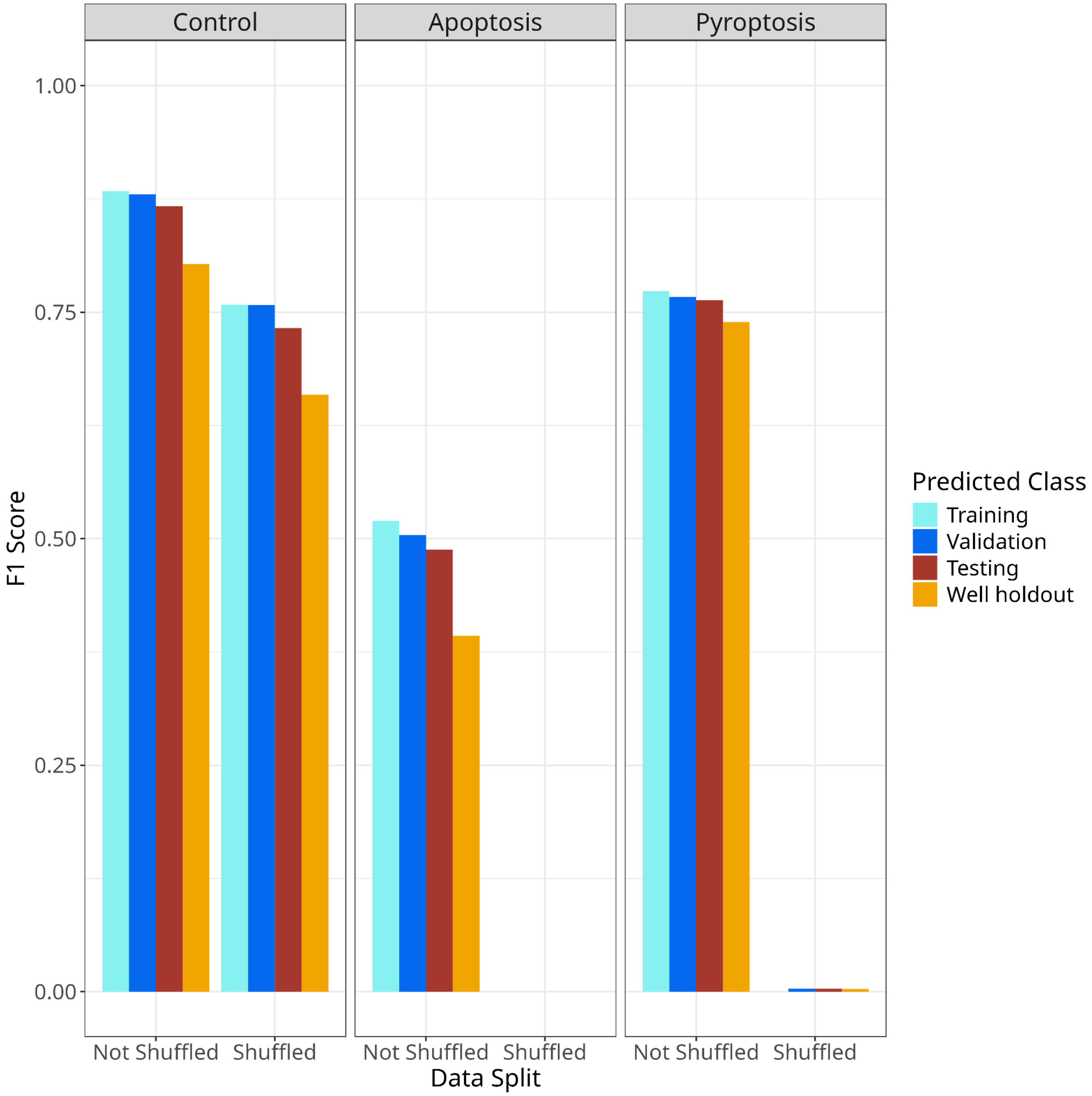

**Supplemental Figure 14:** *F1 scores for predicting cell death across classes, data splits, and data shuffles.*

F1 score is the harmonic mean of the precision and recall of a predicted class.

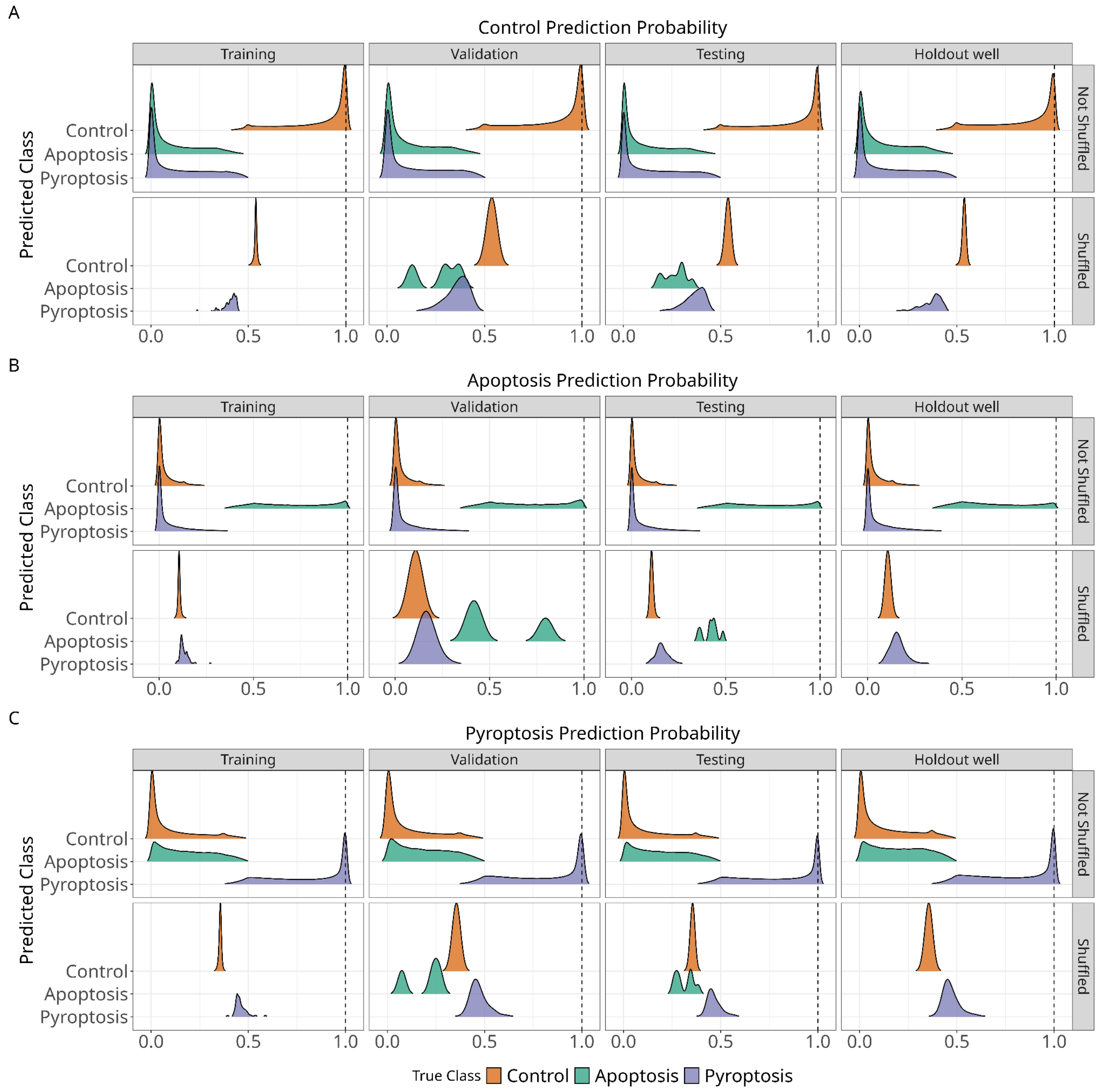

**Supplemental Figure 15:** *Predicted probability distribution of single cells across cell death class, data split, and data shuffle.*

**(A)** Control class true label predicted probabilities. **(B)** Apoptosis class true label predicted probabilities. **(C)** Pyroptosis class true label predicted probabilities. The dotted line represents a predicted probability of 1 for each class and data split. The holdout facet represents both well and treatment holdouts.

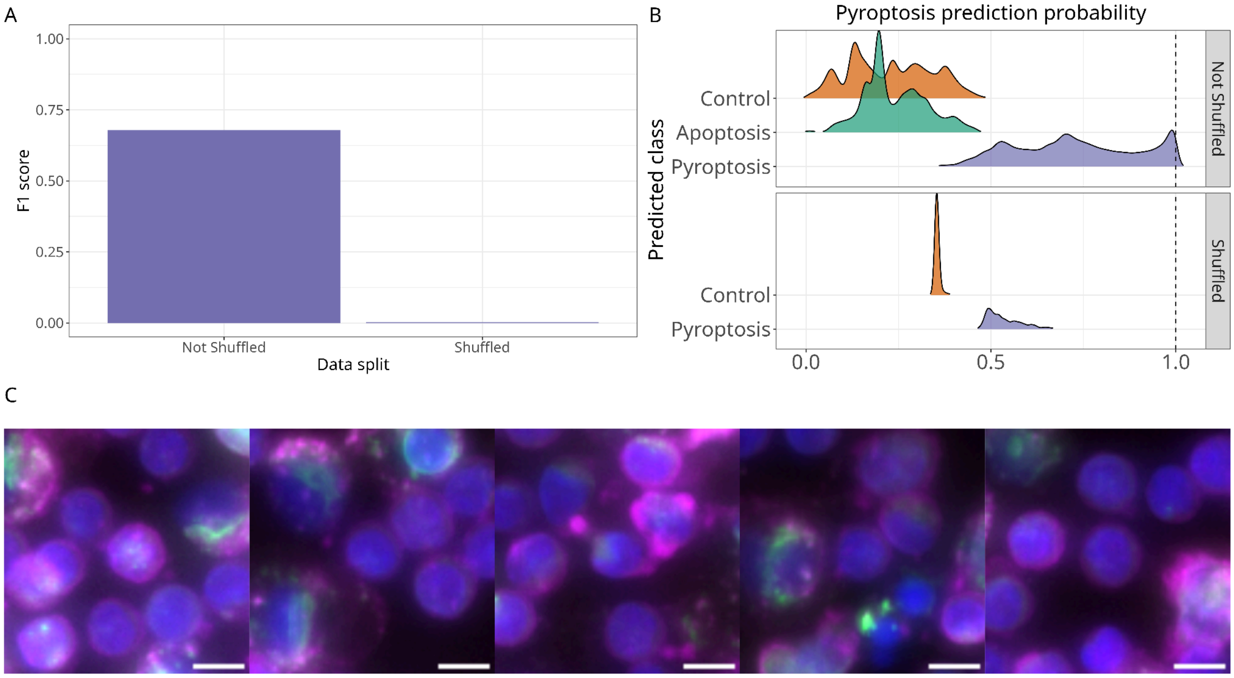

**Supplemental Figure 16:** *Treatment holdout metrics and representations.*

**(A)** The F1 score of the non-shuffled and shuffled model. **(B)** The predicted probability distribution of the classes across the treatment holdout non-shuffled and shuffled model. The apoptosis shuffled density curve is not present as the shuffled model has only zero predictions for the apoptosis class and therefore infinite density at zero (see Figure 3B). **(C)** Representative single cell images treated with a held-out pyroptosis inducer scored with a predicted probability of one. The gasdermin channel (green), actin, golgi, and plasma membrane (magenta) and nuclei (blue) show cleaved Gasdermin D at the cell periphery. Scale bars represent 5 µm.

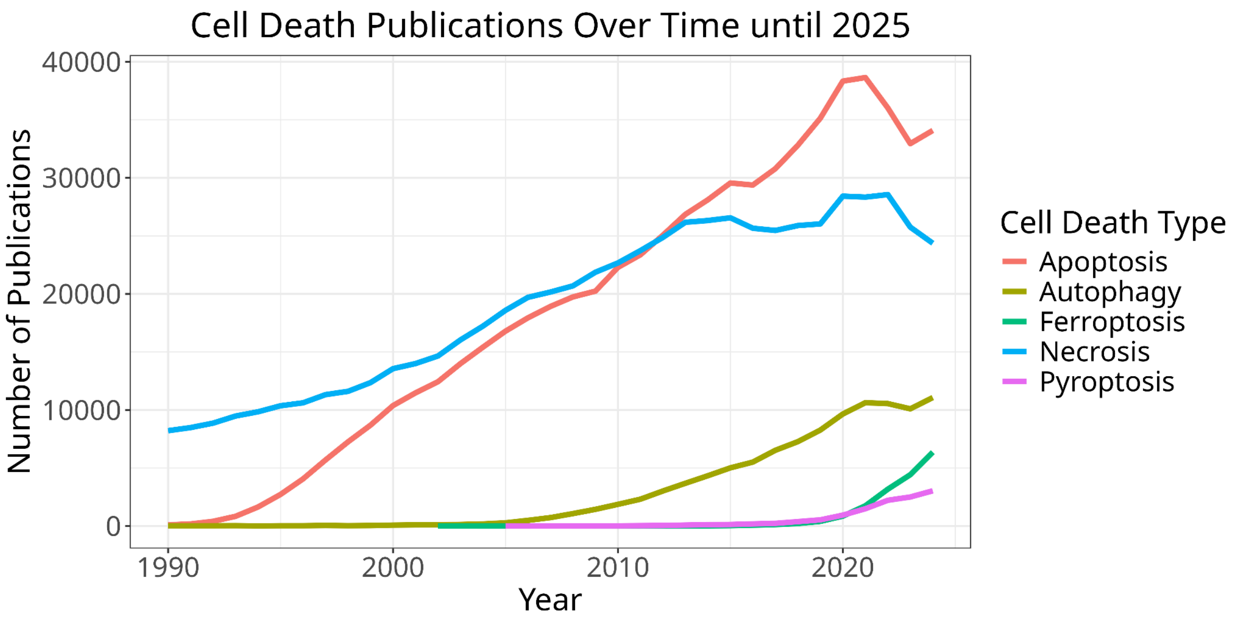

**Supplemental Figure 17:** *The number of publications referencing select forms of cell death over time from 1990 - 2024.*
